# A pontine-specific axonal niche supports de novo gliomagenesis

**DOI:** 10.1101/2024.09.04.611079

**Authors:** Zhigang Xie, Adrija Pathak, Vytas A. Bankaitis

**Affiliations:** Department of Cell Biology & Genetics, Texas A&M University Health Science Center, College Station, TX 77843, USA; Department of Chemistry, Texas A&M University, College Station, TX 77843, USA

## Abstract

Diffuse intrinsic pontine gliomas (DIPGs), a major type of pediatric high-grade gliomas located in the pons, are the leading cause of death in children with brain cancer. A subset (20-25%) of DIPGs harbor a lysine 27-to-methionine (K27M) mutation in *HIST1H3B*, which encodes histone H3.1, and an activating *ACVR1* mutation. The occurrence of this pair of mutations in DIPGs, but not in pediatric gliomas in other anatomical locations, suggests the existence of a pontine-specific niche that favors DIPG gliomagenesis. Unfortunately, the identity of the underlying pontine niche remains elusive as available mouse models fail to recapitulate the anatomic specificity that characterizes DIPGs. Herein we show that the trigeminal root entry zone (TREZ), a pontine structure where several major axon tracts intersect, is enriched with proliferating oligodendrocyte-lineage cells during brainstem development. Introducing both *H3.1K27M* and activating *Acvr1* and *Pik3ca* mutations (which co-occur frequently with *H3.1K27M* in human DIPGs) into the mouse brain leads to rapid gliomagenesis. This pathology recapitulates the pons specificity of DIPGs as glioma cells proliferate on axon tracts at the TREZ. We further show that a hyaluronan receptor important for cell stemness (HMMR) plays a key role in glioma cell proliferation at the TREZ. We propose that *H3.1K27M* and its co-occurring mutations drive pontine specific gliomagenesis by inducing a proliferative response of oligodendrocyte-lineage cells with enhanced stemness on large TREZ axon tracts.

**One-Sentence Summary:** The trigeminal root entry zone underlies pontine-specific gliomagenesis driven by *H3.1K27M* and its co-occurring mutations.

---

Diffuse intrinsic pontine gliomas (DIPGs) are high-grade gliomas located in the pons. DIPGs represent the major cause of brain cancer-related deaths in children. These gliomas occur with an incidence of ∼ 300 new cases each year in the United States alone and a median survival of less than one year (*1, 2*). Over 85% of DIPGs harbor a lysine 27 to methionine (K27M) mutation in genes encoding histone H3.3 or histone H3.1 (*1–6*). These K27M-expressing DIPGs form a major subset of a group of high-grade gliomas termed diffuse midline gliomas – H3K27 altered. At the molecular level, *H3K27M* interferes with the function of polycomb repressive complex 2 which in turn regulates gene expression by catalyzing the mono-, di-, and tri-methylation of H3K27 (*7–10*). Since the identification of *H3K27M* as a major driver mutation in DIPGs, many transplantation-based or genetically engineered animal models have been developed to understand how this mutation cooperates with other DIPG mutations to drive gliomagenesis. Studies of these animal models have revealed several key factors underlying DIPG pathogenesis. These include: (i) stemness of neural stem/progenitor cells (*11–13*), (ii) the developmental program of oligodendrocyte-lineage cells (*14–16*), and (iii) the interaction of neurons with glioma cells (*17–20*). Moreover, studies of these animal models identify several potential molecular targets for treating DIPGs (*17, 21–23*). Despite this major progress, a long-standing puzzle remains unresolved. In human pediatric high-grade gliomas, an obligatory pair of *H3.1K27M* and an activated *ACVR1* mutation are detected exclusively in DIPGs and not in high-grade gliomas in other locations (*3–6*). This anatomical specificity of DIPG mutations strongly implies existence of a pontine-specific niche where the H3.1-K27M/ activated ACVR1 pair support particularly high tumorigenic potential. Unfortunately, this anatomical specificity has neither been recapitulated nor identified in genetically engineered mouse models (*13, 15, 16, 21–23*). Thus, the nature of such a pontine-specific niche defines a major unanswered question in DIPG pathogenesis.

In this study, we provide evidence that the trigeminal root entry zone (TREZ) is a pontine-specific niche that determines the anatomical specificity of the *H3.1K27M*/activated *ACVR1* mutation pair. The TREZ is the ventrolateral pontine region where the trigeminal nerve enters the pons. Several large axon tracts intersect at the TREZ. These include: (i) the trigeminal nerve (which continues as the spinal trigeminal tract within the pons), (ii) the middle cerebellar peduncle, and (iii) the trapezoid body (*24–26*). We show that the TREZ is enriched with proliferating oligodendrocyte-lineage cells during early postnatal brainstem development. Introduction of transposon-based plasmids for combined expression of *H3.1K27M* with activated *Acvr1*/activated *Pik3ca* mutations (which frequently co-occur with *H3.1K27M* in human DIPGs) into the brainstem of mouse embryos during early brain development induce rapid de novo gliomagenesis. This pathology occurs in the ventrolateral pons and is characterized by glioma cell proliferation primarily restricted to the TREZ. This same mutational cohort fails to induce gliomagenesis within the cerebral hemisphere, the thalamus, or the midbrain. Moreover, expression of *H3.3K27M* and its co-occurring mutations, which are detected in both DIPGs and other midline-gliomas in human patients, does not induce TREZ-specific glioma cell proliferation. To elucidate the mechanisms by which glioma cell proliferation is stimulated at the TREZ, we show that HMMR, a hyaluronan receptor important for cell stemness, is critical for the glioma cell proliferation induced by combined expression of *H3.1K27M*, activated *Acvr1*, and activated *Pik3ca*. We propose a model where *H3.1K27M* and its common co-occurring DIPG mutations confer increased stemness to oligodendrocyte-lineage progenitor cells. These cells then persistently proliferate along the large axon tracts in the TREZ.

## Results

### High density and expansion of proliferating oligodendrocyte-lineage cells in the TREZ during development

Given the role of the oligodendrocyte-lineage development program in DIPG pathogenesis (*14–16*), we assessed which regions of the brainstem exhibited particularly active proliferation of oligodendrocyte-lineage cells. The expansion of the oligodendrocyte-lineage cells in the brainstem occurs primarily at early postnatal stages, with the peak of cell proliferation around P4 (*27*). We therefore immunostained brainstem sections of P4 mouse pups. Sagittal sections were used because several hallmark brainstem structures can be identified based on DAPI nuclear staining in these sections (fig. S1). Proliferating oligodendrocyte-lineage cells were identified by co-expression of MKI67 (the most commonly used cell proliferation marker) and OLIG2, a transcription factor specifically expressed in oligodendrocyte-lineage cells (*28, 29*). At this stage, proliferating oligodendrocyte-lineage MKI67^+^ OLIG2^+^ cells were most concentrated in the TREZ relative to other areas of the brainstem (Fig. 1). This profile persisted through P8, although the densities of proliferating oligodendrocyte-lineage cells at the TREZ were diminished compared to those observed at P4 (fig. S2).

**Fig. 1.**
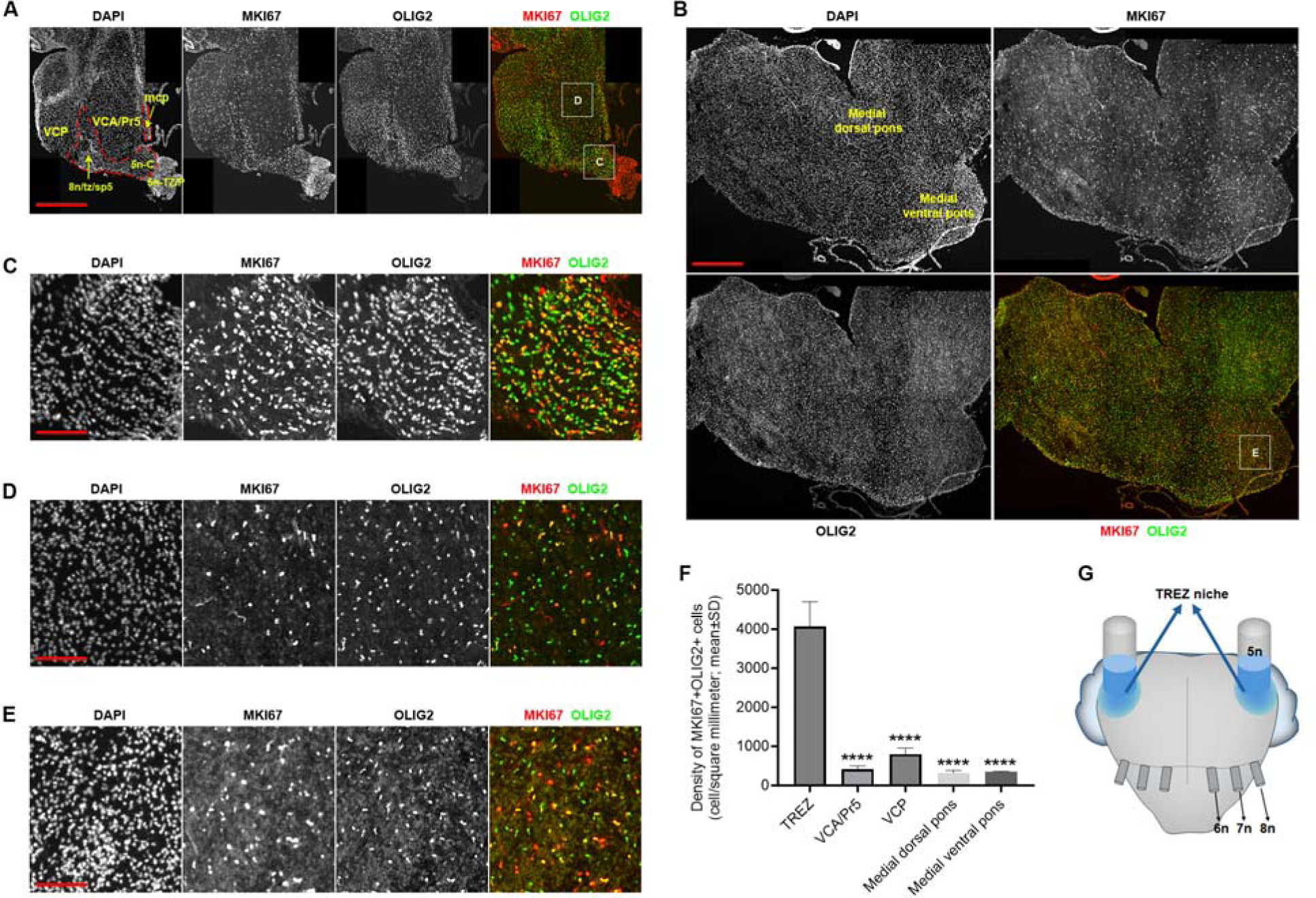
TREZ at the ventrolateral pons is enriched with proliferating oligodendrocyte-lineage cells during early postnatal development. (**A**) Representative composite images showing co-expression of MKI67 and OLIG2 in the lateral mouse brainstem at P4. (**B**) Representative composite images showing co-expression of MKI67 and OLIG2 in the medial mouse brainstem at P4. (**C-E**) Boxed areas in (A) and (B) are shown at higher magnifications to illustrate the high density of proliferating oligodendrocyte-lineage cells (i.e. MKI67^+^OLIG2^+^ cells) at TREZ. Scale bars: 500μm in (A) and (B), 100μm in (C-E). (**F**) Quantification of the density of proliferating oligodendrocyte-lineage cells at different anatomical locations of the brainstem. Quantifications were performed on four embryos for each anatomical location (n=4). ****p<0.0001, one-way ANOVA. (**G**) Schematic representation of the TREZ niche. Ventral view of the brainstem is shown. Anatomical areas of the brainstem are indicated by abbreviations. 5n-c: central portion of the trigeminal nerve; 5n-TZ/P: transitional zone and peripheral portion of the trigeminal nerve; 6n: the 6^th^ cranial nerve; 7n: the 7^th^ cranial nerve; 8n: the 8^th^ cranial nerve; mcp: middle cerebellar peduncle; Pr5: principal sensory trigeminal nucleus; sp5: spinal trigeminal tract; tz: trapezoid body; VCA: ventral cochlear nucleus, anterior; VCP: ventral cochlear nucleus, posterior. The TREZ niche, outlined by dashed lines in (A), includes 5n-c and part of the mcp and 8n/tz/sp5 axon tracts that are close to 5n-c.

### Expression of H3.1K27M and its signature co-occurring mutations leads to rapid de novo gliomagenesis in the TREZ

To determine whether the TREZ was involved in DIPG pathogenesis driven by *H3.1K27M* and activated *Acvr1*, these mutations were deployed to induce de novo gliomagenesis in the mouse pons via in utero electroporation. We designed Tol2 transposon-based plasmids (*30*) for expressing H3.1-K27M and an activated ACVR1 (ACVR1-G356D) (*5*). In addition, as activating mutations of phosphoinositide 3-kinase (PI3K) frequently co-occur with the *H3.1K27M*/activated *ACVR1* in human DIPGs (*3–6*), we also generated a Tol2 transposon-based plasmid for expressing PIK3CA-E545K. This represents an activated form of PIK3CA that is frequently detected in human cancers (*31*). These plasmids, together with a Tol2 transposon-based EGFP plasmid and a conventional plasmid for expressing the Tol2 transposase, were introduced into the brainstem of mouse embryos via in utero electroporation. The transposase catalyzed insertion of the various cassettes for expressing H3.1-K27M, ACVR1-G356D, PIK3CA-E545K, and EGFP (combined expression of these proteins will be referred to as expression of H3.1K27M-AP hereafter) into the genome of transfected cells. EGFP marked the transfected cells and their progenies. In light of the importance of cell stemness in DIPG pathogenesis (*11–13*), these plasmids were introduced into the brainstem at the earliest stage when we were able to reliably perform in utero electroporation (E11.5). We reasoned that, at this stage, the mutations would be expressed in early-stage oligodendrocyte-lineage stem/progenitor cells and would therefore more likely induce or maintain relatively high stemness of the transfected cells. Electroporated embryos were allowed to develop to term, and pontine gliomagenesis was assessed at various postnatal stages. As control (designated as H3.1wt-AP), the same procedure was performed with the exception that the *H3.1K27M* plasmid was replaced with a Tol2 transposon-based plasmid for expressing wild-type H3.1.

A subset of mice born from electroporated embryos displayed hydrocephalus and were promptly euthanized between the stage of weaning (∼ P21) and three months (Table S1). The relatively frequent occurrence of hydrocephalus in both control and experimental groups (Table S1) suggested that this phenotype resulted from the experimental procedure (i.e. in utero electroporation at E11.5). This technical circumstance introduced a variable that complicated using animal survival as an accurate assessment of progression of gliomagenesis. We therefore focused on immunohistological analysis. No tumor cell proliferation was detected in any of the control mice (Fig. 2A). In the H3.1K27M-AP group, however, tumor tissues were observed in the ventrolateral pontine area in mice that did not present symptoms of hydrocephalus (Fig. 2B). Most of these mice developed a head-tilting phenotype at ages ranging from P40 to two months (Table S1 and Movie S1). Immunostaining analysis revealed that proliferating cells expressing H3-K27M (MKI67^+^K27M^+^ cells) were present throughout the lateral pons, with the highest concentration in the TREZ area and the surface of the pons (Fig. 2, C-E). In addition, MKI67^+^K27M^+^ cells appeared to have also spread to medial ventral pons via the middle cerebellar peduncle and the surface of the ventral pons (Fig. 2, C-E). Thus, the distribution of tumor cells within the brainstem corresponded to the TREZ region, although some tumor cells migrated out via axon tracts or exophytic spread. These represented high-grade tumors, as evidenced by extensive expression of ANGPT1 (Fig. 2F), an angiogenesis marker (*32*), and the presence of M-phase K27M^+^ cells (Fig. 2G) within the tumor tissue.

**Fig. 2.**
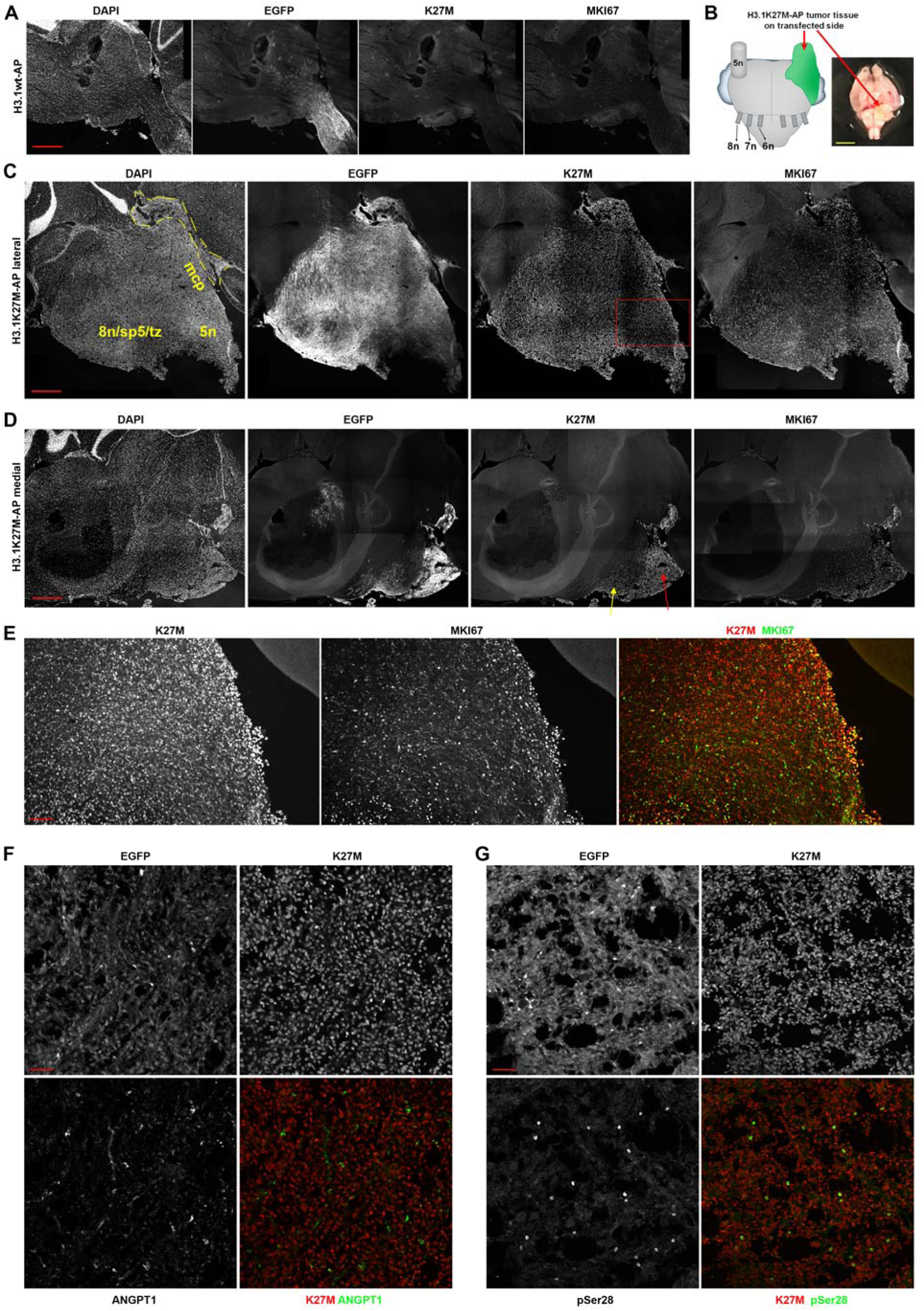
Co-expression of H3.1-K27M and its common co-occurring DIPG mutations leads to de novo formation of high-grade tumors around TREZ. A mixture of transposon-based plasmids for expressing H3.1-K27M, ACVR1-G356D, and PIK3CA (designated as H3.1K27M-AP) was introduced into the brainstem of mouse embryos at E11.5 via in utero electroporation. Mice born from electroporated embryos were analyzed at various postnatal stages. (**A**) Representative composite images of a control experiment (designated as H3.1wt-AP), in which the plasmid for expressing H3.1-K27M was replaced with a plasmid for expressing wild-type H3.1, showing the lack of tumorigenesis and the lack of K27M immunostaining throughout the brainstem. Images are representative of 10 mice at the age of P60-P159. (**B**) Location of the brain tumors induced by H3.1K27M-AP around TREZ. A schematic representation of the tumor location is shown on the left side, and a representative brain sample (P52) with H3.1K27M-AP-induced tumor is shown on the right side. (**C-E**) Images of the brainstem of a P52 mouse in the H3.1K27M-AP group. Composite confocal images of the lateral and medial sagittal sections of the brainstem are shown in (C) and (D), respectively. K27M^+^MKI67^+^ tumor cells were detected throughout the lateral brainstem with the highest density around TREZ and the brainstem surface (outlined by dashed lines) (C). In the medial brainstem (D), K27M^+^MKI67^+^ tumor cells were limited to the surface of the ventral pons (red arrow) and at the medial part of the middle cerebellar peduncle tract (yellow arrow). The boxed area in (C) is shown in (E) at a higher magnification to illustrate that nearly all proliferating (MKI67^+^) cells are K27M^+^. (**F**) An angiogenesis marker, ANGPT1, is expressed extensively in the brain tumors induced by H3.1K27M-AP. (**G**) Mitotic cells are frequently detected in the brain tumors induced by H3.1K27M-AP. Confocal images of (F) and (G) are from a P49 mouse expressing H3.1K27M-AP. Images from (B-G) are representative of nine mice of the H3.1K27M-AP group (P40-P90). 5n: trigeminal nerve; mcp: middle cerebellar peduncle; 8n: the 8^th^ cranial nerve; sp5: spinal trigeminal tract; tz: trapezoid body. Scale bars: 500μm in (A), (C), and (D), 5mm in (B), 100μm in (E), and 50μm in (F) and (G).

Consistent with the human DIPG expression data, and with the involvement of the oligodendrocyte-lineage developmental program (*3–6, 14–16*), the tumor cells induced by H3.1K27M-AP showed enhanced expression of proteins critical for oligodendrocyte-lineage development. These included OLIG2 (Fig. 3, A and B), PDGFRA (Fig. 3C), and ASCL1 (Fig. 3D). Essentially all proliferating tumor cells expressed OLIG2 (Fig. 3, A and B). Furthermore, some cells within the tumor tissues overexpressed NESTIN and GFAP -- markers for neural stem cells and tumor angiogenesis (*33*) and for astrocyte-lineage cells, respectively (fig. S3). These data confirm that the tumor tissues observed in the H3.1K27M-AP group were gliomas and were human DIPG-like with respect to cell lineage developmental programs.

**Fig. 3.**
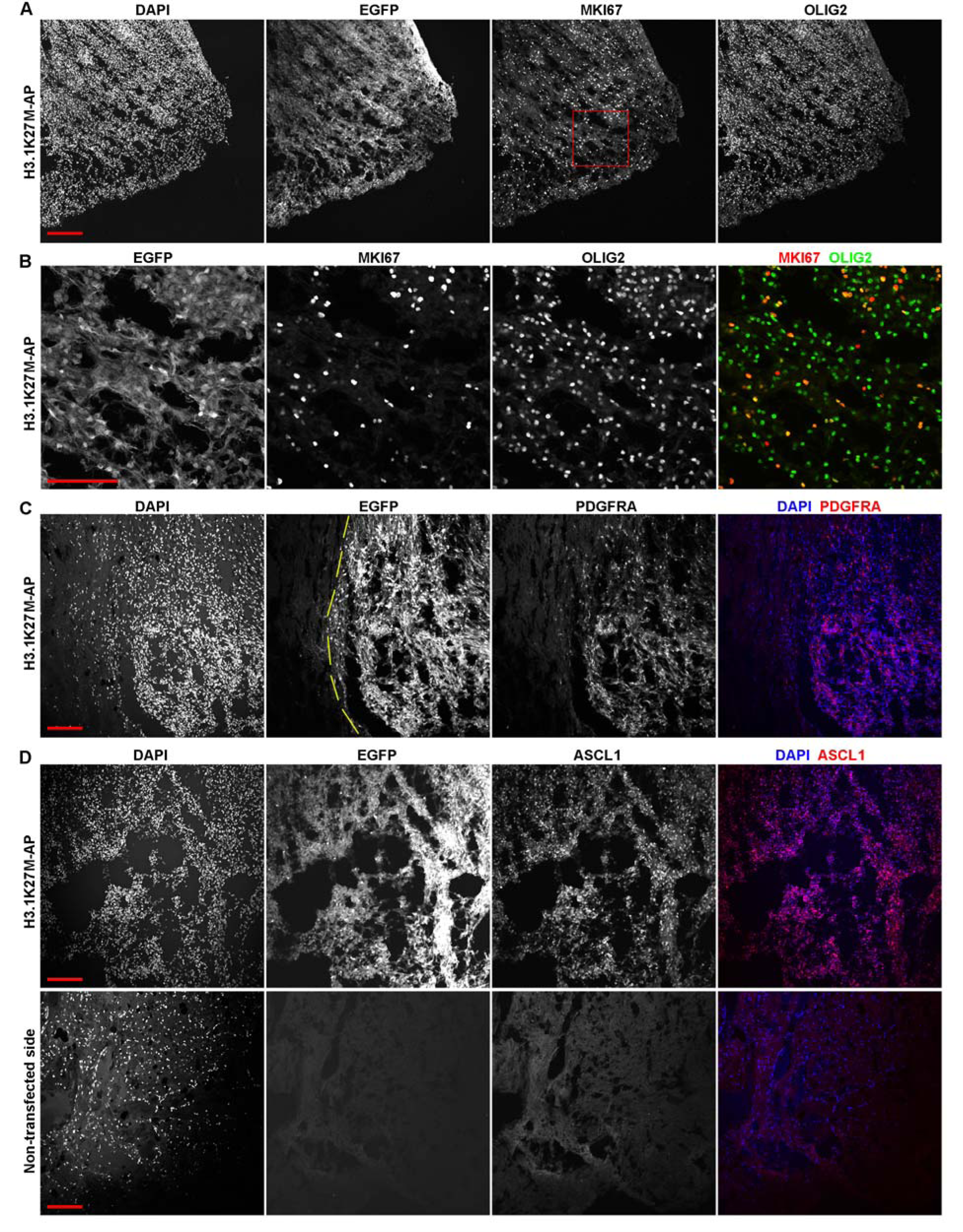
Brain tumors induced by H3.1K27M-AP overexpress oligodendrogenesis-related proteins typical of DIPGs. (**A** and **B**) Confocal images of the brainstem of a P42 mouse expressing H3.1K27M-AP. The boxed area in (A) is shown in (B) at higher magnification to illustrate that nearly all proliferating tumor cells express OLIG2. (**C** and **D**) Confocal images of the brainstem of a P49 mouse showing elevated levels of PDGFRA in tumor cells expressing H3.1K27M-AP compared to nearby non-tumor cells (left side of the dashed lines) (C), or overexpression of ASCL1 in tumor cells compared to non-transfected cells in the contralateral brainstem (D). Scale bars: 100μm.

Interestingly, in a subset of mice expressing H3.1K27M-AP, many of the proliferating K27M^+^ cells did not show EGFP expression (fig. S4, A and B). This observation might result from suppression of EGFP expression (mechanisms unknown), and demonstrated that EGFP expression alone was insufficient for confident identification of all transfected cells and their progenies. Thus, expression of K27M was used in our following analyses for this purpose. While this method could not be used for control groups, where no K27M was expressed, no increased cell proliferation was observed in any brainstem region in the H3.1wt-AP group compared to the group in which EGFP alone was expressed (Fig. 2A and fig. S4, C and D). Thus, tumor cell proliferation observed in the H3.1K27M-AP group is *H3.1K27M*-dependent.

### Glioma cell proliferation induced by H3.1K27M and its co-occurring mutations occurs primarily on axon tracts in the TREZ

In most of the H3.1K27M-AP mice that displayed hydrocephalus, the mice were euthanized before large infiltrative tumors formed. These mice were analyzed for anatomical specificity of glioma cell proliferation induced by H3.1K27M-AP because the dispersion of tumor cells was limited. In addition, we obtained several mice that did not display hydrocephalus and were transfected with H3.1K27M-AP at relatively low efficiencies (by adjusting the parameters of the in utero electroporation procedure). We assessed transfected cell proliferation in various anatomical locations of the brainstem in these mice as well. Strikingly, in all of these mice (ages P20-P91), the TREZ defined the predominant site of cell proliferation (Fig. 4). Compared to other areas, this area had the largest concentration of proliferating cells and the highest fraction of K27M^+^ cells that were proliferating. In areas not directly connected to the TREZ via major axon tracts, few proliferating cells were observed (Fig. 4). While K27M^+^ cells also proliferated on axon tracts away from (but directly connected to) the TREZ, the total numbers of transfected cells in these areas were far less than those at the TREZ (fig. S5). Thus, either these cells were spreading out along axon tracts from the TREZ, or these cells rapidly migrated to the TREZ via axon tracts to continue persistent proliferation, or both. All of these possibilities are consistent with the TREZ representing the predominant site of cell proliferation.

**Fig. 4.**
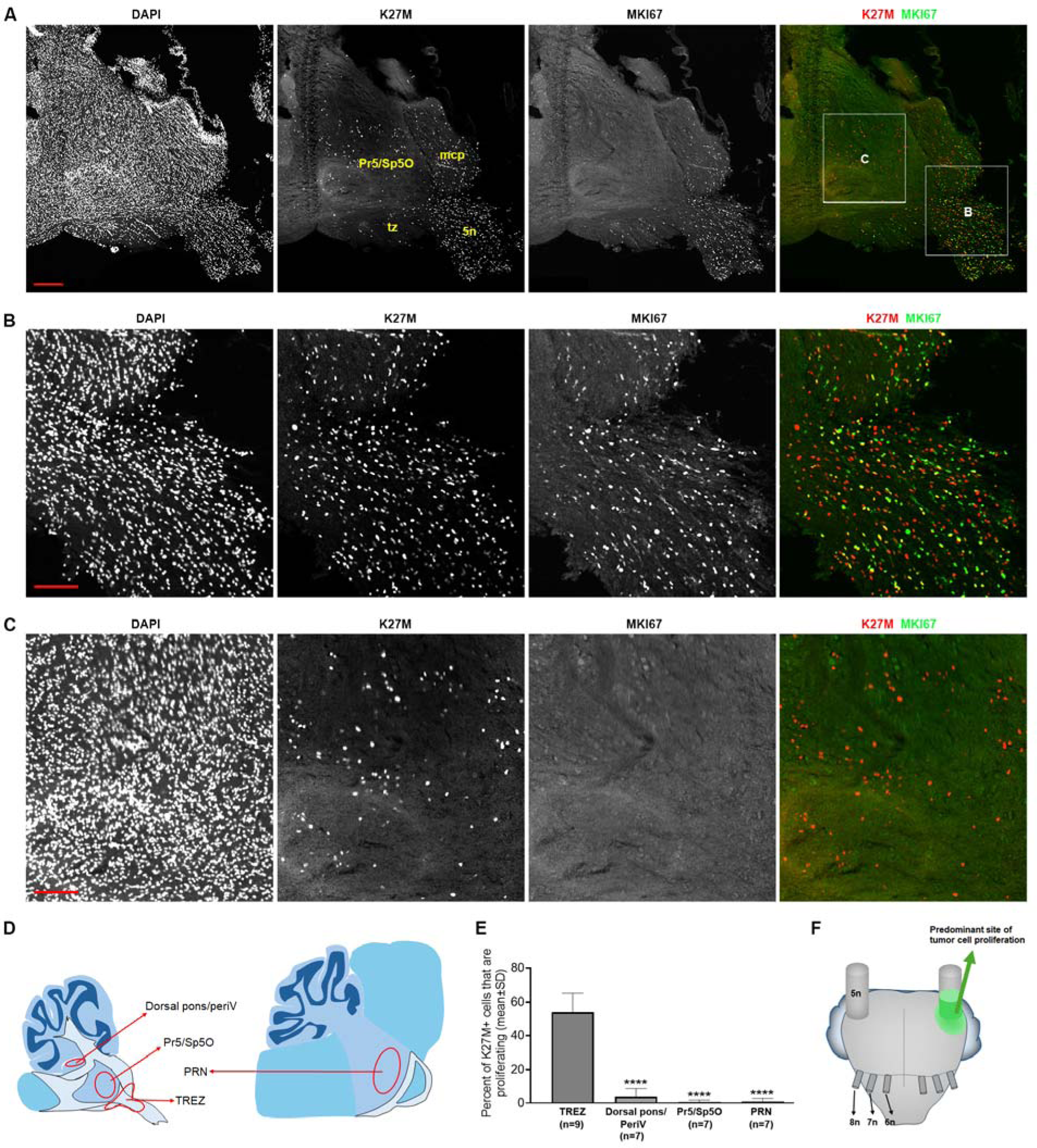
The TREZ is the predominant site of glioma cell proliferation driven by H3.1K27M-AP. (**A-C**) Composite confocal images of the brainstem of a P21 mouse expressing H3.1K27M-AP. Transfected cells were identified by K27M immunoreactivity. Boxed areas in (A) are shown in (B) (TREZ area) and (C) (outside of the TREZ area). Scale bars: 200μm in (A) and 100μm in (B) and (C). (**D** and **E**) Quantification of the proliferation index in transfected cells at different anatomical locations. The schematic in (D) shows the approximate areas used for quantification. ****p<0.0001, one-way ANOVA. The n values indicate the number of embryos used for quantification. (F) A schematic representation showing the predominant site of glioma cell proliferation in the brainstem of mice expressing H3.1K27M-AP. 5n: trigeminal nerve; 6n: the 6^th^ cranial nerve; 7n: the 7^th^ cranial nerve; 8n: the 8^th^ cranial nerve; PeriV: periventricular area; Pr5: principal sensory trigeminal nucleus; PRN: pontine reticular nucleus; Sp5O: spinal trigeminal nucleus, oral part; tz: trapezoid body; TREZ: trigeminal root entry zone.

### H3.1K27M and its co-occurring mutations fail to induce glioma cell proliferation within the parenchyma of the cerebral cortex, in the thalamus, or in the midbrain

As the TREZ is a pontine structure, glioma cell proliferation induced by H3.1K27M-AP in this anatomical space offers a rationale for the specific detection of the *H3.1K27M*/activating *ACVR1* mutation pair in human DIPGs. To further examine this possibility, H3.1K27M-AP mutations were introduced into more rostral parts of the brain at the same embryonic stage (i.e. E11.5). These included the cerebral cortex, the thalamus, and the midbrain. H3.1K27M-AP introduction failed to induce postnatal cell proliferation within the parenchyma of the cerebral cortex, in the thalamus, or in the midbrain (fig. S6, A-D). Interestingly, in a subset of mice (2 of 13 mice with moderate to high transfection efficiencies at the forebrain) periventricular/ventricular tumorigenesis was induced (fig. S6E and Table S1) with most of the proliferating tumor cells expressing OLIG2 (fig. S7). In control mice where *H3.1K27M* was replaced with *H3.1wt*, no such periventricular/ventricular glioma-like cell proliferation was observed (Table S1). The induction of periventricular/ventricular gliomagenesis by H3.1K27M-AP is not inconsistent with the general concept that the pair of *H3.1K27M* and activating *ACVR1* mutation are specific to DIPGs. Rather, these data are consistent with reports that H3K27M-positive periventricular/ventricular gliomas do occasionally occur in human patients (*34–36*). In sum, our demonstration that H3.1K27M-AP failed to induce gliomagenesis in rostral areas of the brain further emphasizes the specific role of the TREZ in H3.1K27M-AP-induced gliomagenesis.

### Expression of H3.3-K27M and its signature co-occurring mutations leads to de novo gliomagenesis in the brainstem outside of the TREZ

We considered the possibility that the TREZ-specificity of H3.1K27M-AP-induced gliomagenesis is driven by technical factors related to how mutations are introduced rather than some fundamental biological foundation. For instance, our electroporation procedure might lead to high transfection efficiencies only in cells destined to populate the TREZ, and therefore selectively drive gliomagenesis from those cells due to their high expression of the mutant proteins. To test this possibility, we used the same approach to induce de novo gliomagenesis with mutational combinations that present different anatomical specificities. To that end, we chose a combination of *H3.3K27M* and its common DIPG co-occurring genetic elements that lead to gliomagenesis both in the pons and in other midline structures in human patients (*3–6*). We reasoned that if the anatomical specificity of H3.1K27M-AP was due to technical factors, then expression of H3.3K27M and its DIPG co-occurring genetic elements would recapitulate the anatomical TREZ-specificity we observed for H3.1K27M-AP-induced gliomagenesis. By contrast, if the anatomical specificity of H3.1K27M-AP-induced gliomagenesis truly reflected a specific role of the TREZ, then expression of H3.3K27M and its DIPG co-occurring genetic elements would not exhibit TREZ-specificity.

In human DIPG patients, the most frequent co-occurring genetic elements with *H3.3K27M* include mutations in *TP53*, overexpression of *PDGFRA*, and overexpression of *CDK/CCND* genes (*3–6*). We therefore designed Tol2 transposon-based plasmids for expressing *H3.3K27M*, *Trp53-R172H* (*37*), wild-type *Pdgfra*, and wild-type *Ccnd2* (hereafter referred to as H3.3K27M-TPC). These plasmids, together with a Tol2 transposon-based EGFP plasmid and a conventional plasmid for expressing the transposase Tol2, were electroporated into the brainstem of mouse embryos at E11.5. Postnatal gliomagenesis in mice born from the electroporated embryos was subsequently analyzed. In mice with moderate transfection efficiencies, no tumor-like cell proliferation was observed in any regions of the brainstem (fig. S8). In mice with high transfection efficiencies, however, extensive tumor-like cell proliferation was observed in the brainstem (fig. S9). No appreciable cell proliferation was induced in the brainstem of control mice (in which H3.3-K27M was replaced with wild-type H3.3) with high transfection efficiencies (fig. S9). These data demonstrated that the tumor-like cell proliferation was *H3.3K27M*-dependent. Immunostaining analysis revealed two types of transfected cell clusters -- one in which only a subset of proliferating cells expressed OLIG2 (fig. S10A), and one in which nearly all proliferating cells expressed OLIG2 (fig. S10B). Intriguingly, whereas extensive glioma-like cell proliferation was observed in various brainstem locations, no such proliferation was recorded at the TREZ. This was the case even though high concentrations of K27M^+^ cells populated the TREZ (fig. S11). While the mechanisms underlying the absence of H3.3K27M-TPC-induced cell proliferation at TREZ remains to be determined, these data argue that the specific role of the TREZ in H3.1K27M-AP-induced pontine gliomagenesis is not an artifact of technical factors related to selective electroporation bias.

### Role of HMMR in H3.1K27M-AP-induced pontine gliomagenesis

Given the cardinal signatures of cell stemness and the oligodendrocyte developmental program in DIPG pathogenesis (*11–16*), our data support a model where H3.1K27M-AP enhances stemness of oligodendrocyte-lineage stem/progenitor cells. This enhancement is in turn amplified by persistent proliferation in the uniquely supportive TREZ large axon tract niche. This concept predicts that interfering with cell stemness, and/or the interaction of oligodendrocyte-lineage cells with axons, will inhibit H3.1K27M-AP-induced gliomagenesis. In this regard, we interrogated the role of HMMR, a multifunctional protein important for cell migration, brain development, and gliomagenesis (*38, 39*). In mice, the *Hmmr* transcript is highly expressed in the developing central nervous system (NCBI gene ID#15366). HMMR not only plays a key role in cell stemness (*40, 41*), but also serves as a receptor for hyaluronan -- an extracellular matrix proteoglycan important for the proliferation and differentiation of oligodendrocyte-lineage cells during axonal myelination (*42*). Indeed, we observed increased expression of HMMR in the TREZ niche during mouse brainstem development (Fig. 5A). Moreover, proliferating glioma cells transfected with H3.1K27M-AP exhibited elevated levels of HMMR both before (Fig. 5B) and after (fig. S12) large tumor formation.

**Fig. 5.**
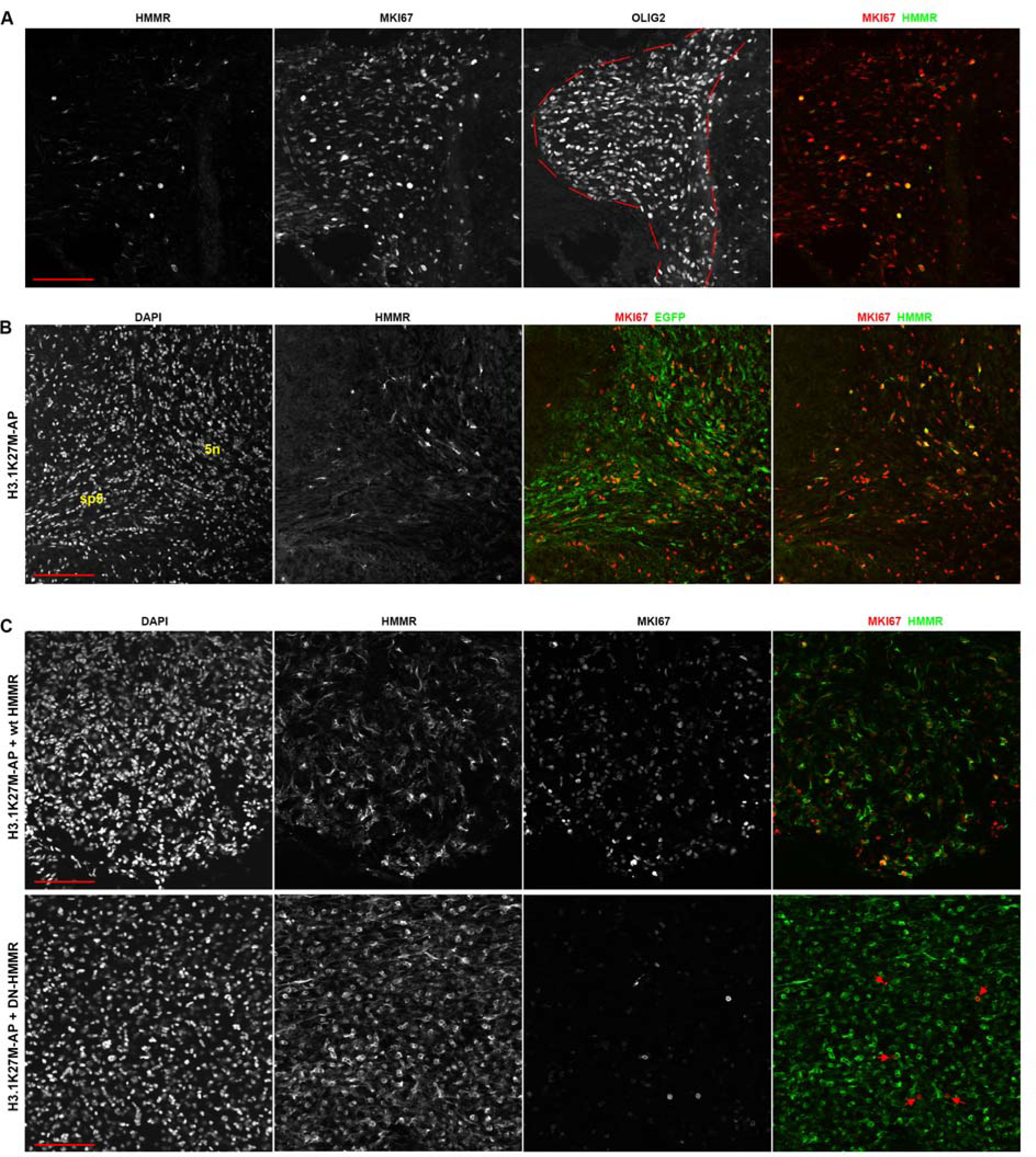
Interfering with HMMR function impairs glioma cell proliferation induced by H3.1K27M-AP. (**A**) Expression of HMMR in TREZ at P4. HMMR expression levels are higher in TREZ (outlined by dashed lines) compared to nearby regions. Images are representative of four embryos. (**B**) Confocal images of the brainstem of a P22 mouse showing elevated expression of HMMR in proliferating glioma cells. The approximate regions of the axon tracts at TREZ are indicated in the DAPI panel. Images are representative of all of the mice (nine of the H3.1K27M-AP group with moderate number of proliferating glioma cells and no large tumor bulk) that were subjected to HMMR immunostaining analysis. (**C**) A dominant-negative HMMR (DN-HMMR) inhibits glioma cell proliferation induced by H3.1K27M-AP. Confocal images from a P60 control mouse (H3.1K27M-AP co-transfected with wild-type HMMR) and a P55 DN-HMMR mouse (H3.1K27M-AP co-transfected with DN-HMMR) are shown. HMMR immunoreactivity was used to identify transfected cells. Co-expression of DN-HMMR, but not wild-type HMMR, markedly reduced the proliferation of cells transfected with H3.1K27M-AP. Note that proliferating cells in the DN-HMMR group (indicated by arrows) generally exhibited low levels of HMMR immunoreactivity. Images are representative of three control mice (P28-P61) and seven DN-HMMR mice (P21-P60). 5n: trigeminal nerve; sp5: spinal trigeminal tract. Scale bars: 100μm.

To determine whether HMMR supports pontine H3.1K27M-AP-induced gliomagenesis, a Tol2 transposon-based plasmid for expressing a mutant HMMR (DN-HMMR) was co-electroporated with H3.1K27M-AP expression plasmids into the brainstem of E11.5 mouse embryos. DN-HMMR harbors several C-terminal amino acid substitutions that abrogate hyaluronan binding, and its expression exerts dominant-negative effects on hyaluronan signaling (*43*). Gliomagenesis in the brainstem of mice born from the electroporated embryos was subsequently analyzed. In the control group where DN-HMMR was replaced by wild-type HMMR, nearly all cells with high HMMR immunoreactivity were proliferative (Fig. 5C). This pattern was indistinguishable from that recorded for mice transfected with H3.1K27M-AP alone (Fig. 5B and fig. S12). In the DN-HMMR group, however, the majority of cells with high HMMR immunoreactivity were not proliferative (Fig. 5C). While gliomagenesis still occurred, cell proliferation was restricted to those cells with low HMMR immunoreactivity (Fig. 5C, Table S1) -- i.e. those cells exhibiting low ectopic expression of DN-HMMR. These data indicate that interfering with HHMR activity inhibits pontine H3.1K27M-AP-induced glioma cell proliferation.

## Discussion

In this study, we identify the TREZ as a pontine-specific region that uniquely supports gliomagenesis induced by *H3.1K27M* and its common co-occurring DIPG mutations. As such, this study provides an answer to the long-standing puzzle why the *H3.1K27M*/activating *ACVR1* mutation pair are cardinal features of pediatric DIPGs, but are not featured in pediatric high-grade gliomas that arise in other anatomical locations. The case is as follows. First, we show that co-expression of *H3.1K27M*, *Acvr1-G356D*, and *Pikca3-E545K* (i.e. H3.1K27M-AP) induces rapid de novo gliomagenesis in the mouse pons resulting from glioma cell proliferation specifically at the TREZ. H3.1K27M-AP failed to induce gliomagenesis in other anatomical areas of the brain. Second, co-expression of *H3.3K27M* and its common co-occurring DIPG mutations (i.e. H3.3K27M-TPC), a combination that does not display pontine-specificity in human patients, induced de novo gliomagenesis in the brainstem outside of the TREZ but not in the TREZ itself. Finally, interfering with activity of HMMR, a hyaluronan receptor important for cell stemness, markedly impaired H3.1K27M-AP-induced glioma cell proliferation. We interpret the collective data in a model where: (i) H3.1K27M-AP enhances oligodendrocyte-lineage stem/progenitor cell stemness, (ii) the large axon tracts of the TREZ provide a uniquely supportive niche for persistent proliferation of these stem-like cells, and (iii) HMMR activity is a central component of the signaling pathway(s) that drive persistent proliferation of the stem-like cells. In that regard, we note our collective results are consistent with involvement of the trigeminal nerve in a subset of DIPG patients (*44*).

Previous studies employed genetically engineered mouse models that failed to reproduce the anatomical specificities of genetic mutations in pediatric high-grade gliomas (*13, 15, 16, 21, 22*). In several of these studies, co-expression of *H3.1K27M* and activating mutations in the *Acvr1* and *Pik3ca* pathways were used (*16, 21, 23*). However, these studies either did not assess whether (or how) these mutations led to pontine specific gliomagenesis, or reported gliomagenesis in anatomical regions inconsistent with those of human DIPG patients. A major distinction between our study and the previous ones is the embryonic stages at which we introduced the gliomagenic mutations. In the previous studies, the mutations were introduced or expressed at E12.5, at the neonatal stage, or in OLIG2^+^ progenitors and their progeny (i.e. not in earlier stage stem/progenitor cells that do not express OLIG2) (*16, 21, 23*). By contrast, our system introduced the genetic elements into the brainstem at E11.5. At this stage, most brain regions (including the brainstem) are composed of a thin layer of neuroepithelium (*45*), which is populated by early-stage neural stem cells that will ultimately give rise to both neurons and glial cells. We believe this experimental difference to be highly significant as cell stemness is a critical factor in DIPG pathogenesis (*11–13*). We posit that rapid gliomagenesis is induced when the H3.1K27M-AP mutations hit early-stage stem/progenitor cells of the oligodendrocyte-lineage exposed to the TREZ microenvironment. We believe this is the underlying reason for why our data diverge from those of previous studies using similar (but not identical) genetically engineered mouse models. We further suggest that mortality as a result of the rampant H3.1K27M-AP-induced gliomagenesis in the TREZ would precede development of gliomagenesis in other anatomical locations of the brain.

Why is the TREZ uniquely supportive for H3.1K27M-AP-induced gliomagenesis compared to other brain regions? The TREZ is anatomically unique in that several large, myelinated axon tracts intersect at this area. These include: (i) the trigeminal nerve -- the largest facial nerve that continues as the spinal trigeminal tract within the pons, (ii) the middle cerebellar peduncle -- a prominent axon tract that connects the pontine nucleus with the contralateral cerebellum as part of the cortico-ponto-cerebellar pathway, and (iii) the trapezoid body -- a major axon tract that connects the cochlear nucleus with the superior olivary complex as part of the auditory pathway (*24–26*). Given that myelination of axons is the central function of oligodendrocytes, stimulatory effects from these large axon tracts likely drive the persistent proliferation of oligodendrocyte-lineage stem/progenitor cells whose stemness is enhanced by H3.1K27M-AP. Our data on HMMR are consistent with this possibility. HMMR promotes stemness (*40, 41*), and is a receptor for the important extracellular matrix glycosaminoglycan hyaluronan that mediates the interaction between neurons and oligodendrocyte-lineage cells during myelination (*42*). Importantly, expression of a hyaluronan-binding-deficient HMMR does not support H3.1K27M-AP-induced glioma cell proliferation (Fig. 5). While HMMR is also implicated in mitotic spindle positioning and cell cycle progression (*38, 39*), the defects in gliomagenesis that we observed are unlikely to be indirect effects of simple defects in cell cycle progression. Analyses of two different lines of *Hmmr* knockout mice revealed that absence of HMMR or deletion of HMMR C-terminus compromised neural stem cell self-renewal but not cell cycle progression per se in the brain (*46, 47*). In this study, we used a mutant HMMR harboring several C-terminus mutations that abolish hyaluronan binding (*43*). Thus, we believe it more likely that HMMR signaling supports H3.1K27M-AP-induced gliomagenesis by promoting cell stemness and hyaluronan-dependent interaction between neurons and oligodendrocyte-lineage cells.

The particularly important role of the TREZ in H3.1K27M-AP-induced pontine gliomagenesis is consistent with recent findings that neuron-glioma cell interactions are key factors stimulating gliomagenesis (*18–20*). We also establish this point in our study. We emphasize that, in contrast to H3.1K27M-AP-induced gliomagenesis, H3.3K27M-TPC-induced glioma cell proliferation was limited to anatomical areas outside of the TREZ (fig. S11). This observation suggests that, although the TREZ may play important roles in both H3.1K27M-dependent and in H3.3K27M-dependent pontine gliomagenesis, the precise roles are likely different in these two contexts. Thus, elucidating biochemical nature of the TREZ that makes it particularly supportive of *H3K27M*-dependent gliomagenesis is a direction that promises fundamental clues to the molecular basis of DIPG pathogenesis.

## Acknowledgments

Confocal images are acquired on a Nikon confocal microscope maintained by the Department of Cell Biology and Genetics at Texas A&M University Health Science Center.

## Author contributions

Conceptualization, Methodology, and Visualization: ZX. Investigation: ZX, AP. Funding acquisition: ZX, VAB. Project administration: ZX. Supervision: ZX, VAB. Writing – original draft: ZX. Writing – review & editing: ZX, VAB.

## Data and materials availability

All data are available in the main text or the supplementary materials. All plasmids generated in this study will be available upon reasonable request.

## Supplementary Materials

Materials and Methods

Figs. S1 to S12

Table S1

Movie S1

