## Supplementary Materials for "A pontine-specific axonal niche supports de novo gliomagenesis"

**The PDF file includes:**

Materials and Methods

Figs. S1 to S12

**Other Supplementary Materials for this manuscript include the following:**

Table S1

Movie S1

Materials and Methods

Plasmids

Transposon-based plasmids for expressing wild-type or mutant proteins were generated using the Tol2 transposon system (*30*). A 4.0kb fragment carrying the coding sequence for Cre:GFP flanked with a CAG promoter and a rabbit globin polyA sequence was cut from pCAG-Cre:GFP (*48*) with SpeI/HindIII and inserted into the XbaI/HindIII sites of pMiniTol2 (*30*). The 1.8kb EcoRI/NotI fragment of the insert containing the Cre:GFP coding sequence was then replaced with a 0.7kb EcoRI/NotI fragment containing EGFP coding sequence obtained via polymerase chain reaction (PCR) using pEGFPc1 (Clontech) as the template. In the resulting plasmid (referred to as pMiniT2-EGFP), the EGFP expression cassette (CAG promoter, EGFP coding sequence, and rabbit globin polyA sequence) was flanked by Tol2 inverted terminal repeats (ITRs).

To construct transposon plasmids for expressing glioma-related proteins, coding sequences for these proteins were amplified via reverse transcription (RT)-PCR using complementary DNA (cDNA) prepared from an E14.5 mouse embryonic brain, and used to replace the EGFP coding sequence of pMiniT2-EGFP. These sequences include *Hist1h3i* (encoding H3.1), *H3f3a* (encoding H3.3), *Acvr1*, *Pik3ca* (encoding the catalytic subunit of phosphoinositide-3-kinase), *Trp53*, *Pdgfra*, *Ccnd2*, and *Hmmr*. Note that, while in humans both *HIST13HB* and *HIST1H3I* encode H3.1, in mice only *Hist1h3i* encodes H3.1 (mouse *Hist1h3b* encodes H3.2). For expressing mutant proteins H3.1-K27M, H3.3-K27M, ACVR1-G356D, PIK3CA-E545K, TRP53-R172H, and DN-HMMR (which harbors the mutations K723E, K727E, K748N, R749W, and K750E), site-directed mutagenesis was performed to introduce the corresponding mutation into the coding sequences. Both wild-type and mutant coding sequences were verified by sequencing.

Animals and in utero electroporation

Mice were on the background of C57BL6, and were maintained in the animal facility of Texas A&M University Health Science Center. All animal work was performed according to protocols approved by the Institutional Animal Care and Use Committee (IACUC) of Texas A&M University and the animal use guidelines of the National Institute of Health. The in utero electroporation procedure was performed similar to that described in our previous studies with several modifications (e.g. on the site of DNA injection and the orientation of the electrodes) (*49-51*). Briefly, mouse breeding was set up to produce embryos at the stage of E11.5. The date of plug observation was designated as E0.5. At the stage of E11.5, the pregnant mouse was anesthetized and placed onto a surgical platform. An abdominal incision at the size of about 1 centimeter was made, and the uteri were gently pulled out of the abdominal cavity. For brainstem transfection, a mixture of the DNA solution (prepared in sterile water, mixed with ~0.05% Fast Green dye) was injected into the fourth ventricle via a sterile glass micropipette, which were inserted into the aqueduct with the tip pointing toward the fourth ventricle. This method of injection ensured that the brainstem area to be transfected would not be disrupted mechanically by micropipette insertion. Following DNA solution injection, five electric pulses (~17 volts; 50msec on/950msec off; generated from an ECM830 square wave electroporator) were delivered across the mouse embryo via tweezer-like disc electrodes. During the delivery of the electric pulses, the negative electrode disc was positioned on the back of the head of the mouse embryo (the direct contact was on the uterine wall), and the positive electrode disc was positioned at the bottom of the embryo. By this positioning, the electric pulses were expected to be delivered across the pontine flexure. For forebrain/midbrain electroporation, the DNA solution was injected from the right lateral ventricle into the lateral ventricles, the third ventricle, and the aqueduct. After DNA injection, the electrode discs were placed on opposite sides of the embryonic head so that delivery of the electric pulses would result in transfection of cells at the cerebral cortex (where cells took up the plasmids from the lateral ventricle), the thalamus (where cells took up the plasmids from the third ventricle), and the midbrain (where cells took up the plasmids from the aqueduct). After the delivery of the electric pulses, the uteri were placed back into the original position in the abdominal cavity of the dam, the incision was sutured, and the dam was placed in a warm location for recovery. Aseptic handling, analgesics (pre- and post- operation), and post-operational monitoring were used according to the approved methods by the IACUC of Texas A&M University. The electroporated embryos were allowed to develop to term, and euthanized at different postnatal or adult stages.

Brain sample collection and processing

A subset of mice born from electroporated embryos developed hydrocephalus at different postnatal stages. These mice were promptly euthanized for brain harvest when the sign of hydrocephalus (i.e. domed head) was observed. A subset of mice expressing H3.1K27M-AP displayed head-tilting phenotype. These mice were also promptly euthanized for brain harvest upon occurrence of the head-tilting phenotype. A subset of mice were also euthanized for brain harvest without the occurrence of any symptoms. For brainstem transfected mice, the whole brain was dissected out upon euthanasia, and divided into two halves by a sagittal midline cut. The forebrain hemispheres were then removed, and the remaining part of the samples (including the brainstem and the cerebellum) were examined for EGFP fluorescence under an epifluorescence inverted microscope (Nikon Eclipse TS100) with a 10x objective. To minimize the effects of transfection locations (e.g. failure to transfect a particular brainstem region) on the specificity of anatomical locations of gliomagenesis, only the samples with clearly visible green fluorescence on both the medial side of the brainstem and the lateral side of the brainstem were collected for further processing and analysis.

For forebrain/midbrain transfected mice, the whole brain was dissected out upon euthanasia, and divided into two halves by a sagittal midline cut. Both halves of the samples were checked for EGFP fluorescence under an epifluorescence inverted microscope (Nikon Eclipse TS100) with a 10x objective. The sample halves with clearly visible green fluorescence on the cerebral cortex and/or the thalamus/midbrain areas were collected for further processing and analysis.

The collected samples were fixed in 2% paraformaldehyde prepared in phosphate buffered saline (PBS) at room temperature for 45 min, soaked in 20% sucrose prepared in PBS until the samples sank, and embedded in Tissue-Tek O.C.T. The embedded samples were stored at -20°C until cryosection preparation. Sagittal sections (20μm) were prepared for brainstem transfected samples, and either sagittal or coronal sections (20μm) were prepared for forebrain/midbrain transfected samples.

Antibodies and fluorescent immunostaining

The following primary antibodies were used for immunostaining analysis. GFP (chicken, Aves Labs, GFP-1020, 1:1000), GFP (goat, Novus Biologicals, NB100-1770, 1:1000), MKI67 (rabbit, Abcam, ab15580, 1:1000), MKI67 (rat, Invitrogen, 14-5698-82, 1:1000), OLIG2 (goat, R&D Systems, AF2418, 1:1000), H3-K27M (rabbit, Abcam, ab190631), HMMR (rabbit, Abcam, ab124729, 1:300), PDGFRA (rabbit, Cell Signaling Technology, 3174S, 1:1000), GFAP (chicken, Aves Labs, GFAP, 1:1000), NESTIN (rat, Abcam, ab81462, 1:1000), ASCL1 (rabbit, Abcam, ab211327, 1:1000), Histone H3 phospho-Ser28 (rat, Abcam, ab10543, 1:1000). The cryosections were incubated in the primary antibodies prepared in PBS containing 3% bovine serum albumin (BSA) and 0.2% Triton-X-100 at room temperature overnight, secondary antibodies prepared in PBS containing 3% BSA and 0.2% Triton-X-100 at room temperature for one hour, and DAPI solution (5μg/ml in PBS) for 5min. All of the secondary antibodies (Jackson Immunoresearch) were conjugated with cyanine dyes and were with minimal cross reactivity. After staining, EverBrite hardset mounting medium (Biotium) was used to mount coverslips.

Confocal microscopy and imaging processing

Images of immunostained brain sections were acquired on a Nikon confocal microscope controlled by the NIS-Elements software. For some of the analyses, confocal images of different regions of the same section were stitched together using the ImageJ software.

Identification of anatomical locations in the mouse brain

The anatomical locations of the brainstem were identified by comparing the pattern of DAPI staining to the pattern of Nissl staining and acetylcholinesterase staining in published mouse brain atlases (*25,26*). Several major axon tracts in the brainstem (such as the trigeminal nerve, the spinal trigeminal tract, the middle cerebellar peduncle, the inferior cerebellar peduncle, and the trapezoid body) revealed by Nissl staining and acetylcholinesterase staining in the brain atlases (*25,26*) can be identified in the corresponding regions within the DAPI stained sections based on the special arrangement of the DAPI-labeled cell nuclei - the cell nuclei on axon tracts are arranged as aligned strings.

Statistical analysis

Statistical analyses were performed using the GraphPad Prism (version 10) software. One-way ANOVA was used to determine the significance of the differences among different groups.

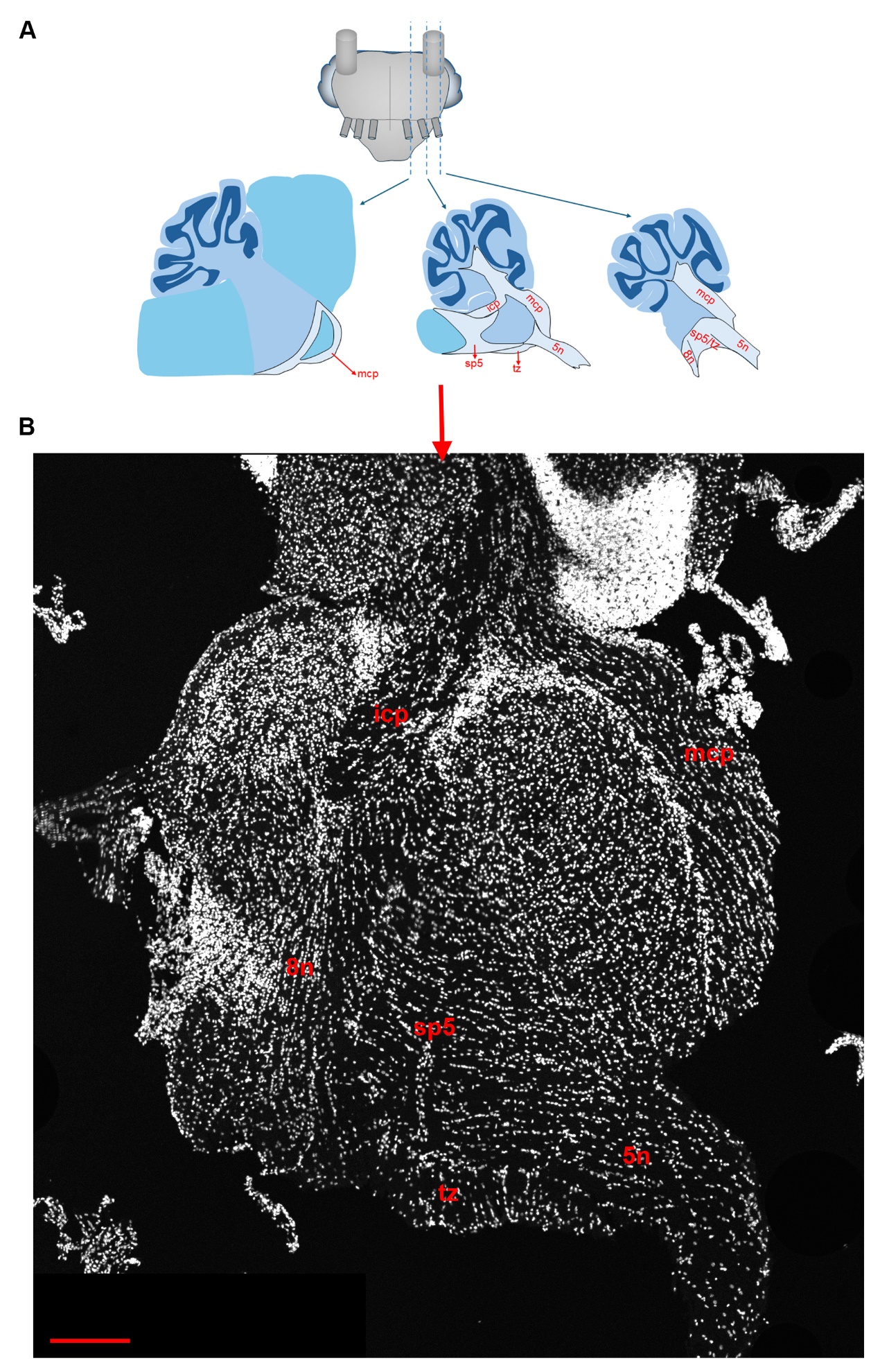

Fig. S1. Identification of anatomical locations in the mouse brainstem based on DAPI staining. (A) Schematic illustrating the anatomical locations in a sagittal section of the mouse brainstem. (B) Composite confocal images showing a DAPI stained lateral brainstem sagittal section prepared from a P20 mouse. Large axon tracts are characterized by nuclei arranged in strings that are generally aligned. These nuclei belong to oligodendrocytes that form the myelin sheath around the axons. The axon tracts identified based on the arrangement of DAPI-labeled nuclei are consistent with those identified based on Nissl staining and acetylcholinesterase staining (*25, 26*). 5n: trigeminal nerve; 8n: the 8^th^ cranial nerve; icp: inferior cerebellar peduncle; mcp: middle cerebellar peduncle; sp5: spinal trigeminal tract; tz: trapezoid body. Scale bar: 200μm.

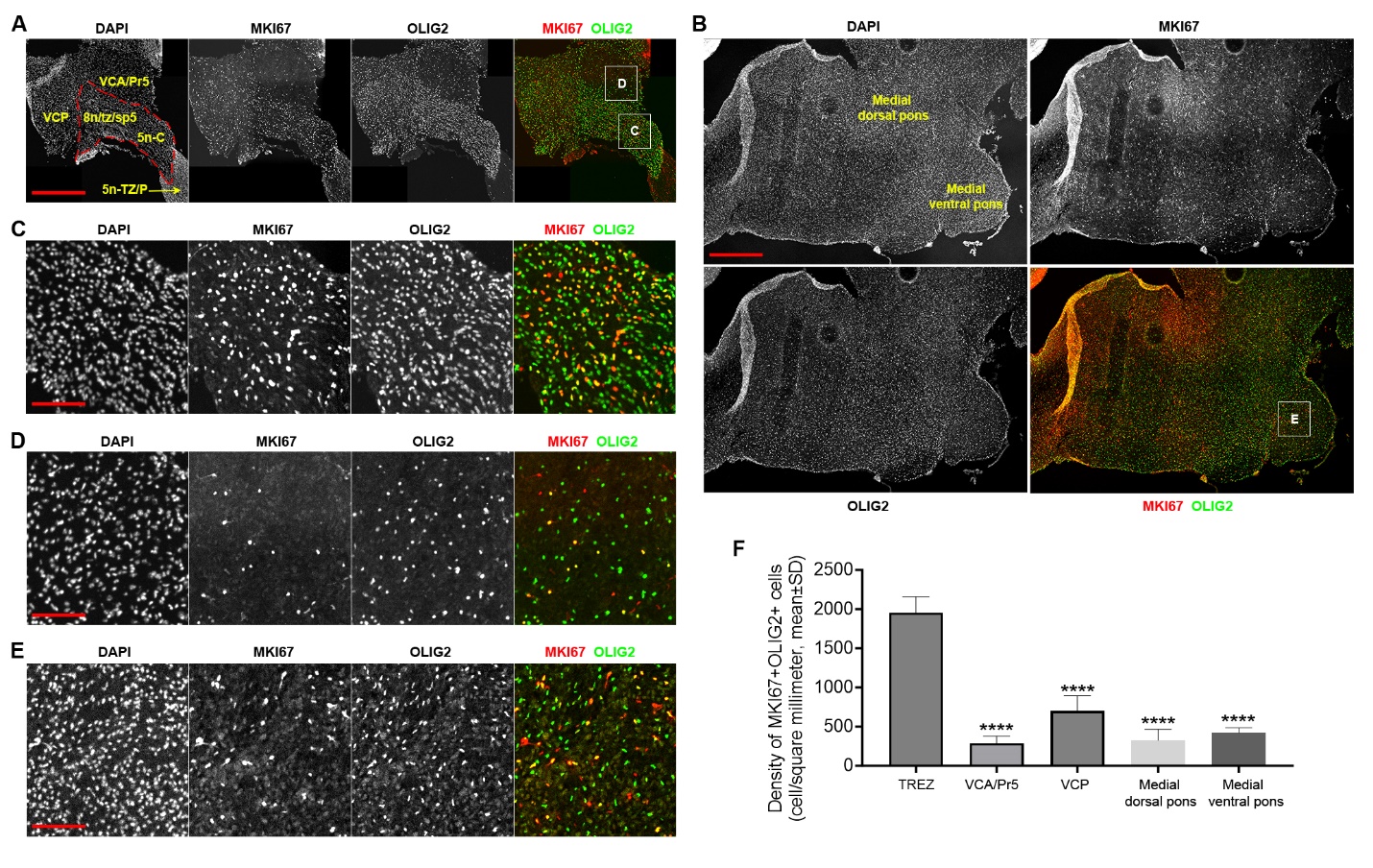

**Fig. S2. TREZ is enriched with proliferating oligodendrocyte-lineage cells at P8.** (**A** and **B**) Representative composite confocal images showing co-expression of MKI67 and OLIG2 in the lateral (A) and medial (B) mouse brainstem at P8. (**C-E**) Boxed areas in (A) and (B) are shown at higher magnifications to illustrate the high density of proliferating oligodendrocyte-lineage cells (i.e. MKI67^+^OLIG2^+^ cells) at TREZ. Scale bars: 500μm in (A) and (B), 100μm in (C-E). (**F**) Quantification of the density of proliferating oligodendrocyte-lineage cells at different anatomical locations of the brainstem. Quantifications were performed on four embryos for each anatomical location (n=4). ****p<0.0001, one-way ANOVA. 5n-c: central portion of the trigeminal nerve; 5n-TZ/P: transitional zone and peripheral portion of the trigeminal nerve; 8n: the 8^th^ cranial nerve; Pr5: principal sensory trigeminal nucleus; sp5: spinal trigeminal tract; tz: trapezoid body; VCA: ventral cochlear nucleus, anterior; VCP: ventral cochlear nucleus, posterior. The TREZ niche is outlined by dashed lines in (A).

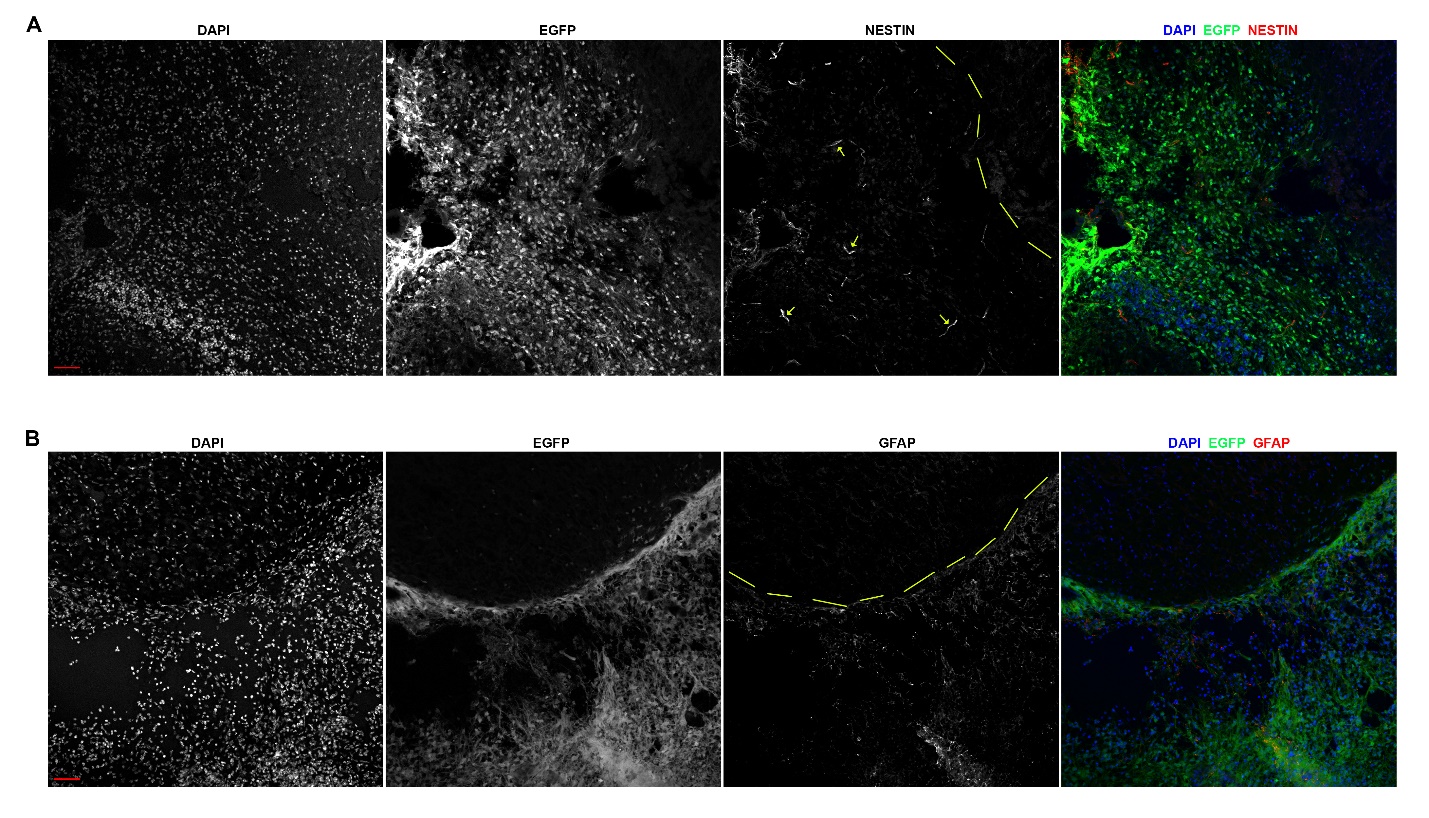

Fig. S3. Increased expression of NESTIN and GFAP in tumor tissues induced by H3.1K27M-AP. Confocal images of the tumor tissue expressing H3.1K27M-AP from a P40 mouse (A) and P64 mouse (B) showing expression of NESTIN and GFAP. Dashed lines approximately outline the edge of the tumor tissue. Levels of NESTIN and GFAP in tumor cells are higher compared to nearby non-tumor cells. Images are representative of nine mice with H3.1K27M-AP-induced pontine gliomas. Arrows in (A) indicate blood vessel-like staining. Scale bars: 100μm.

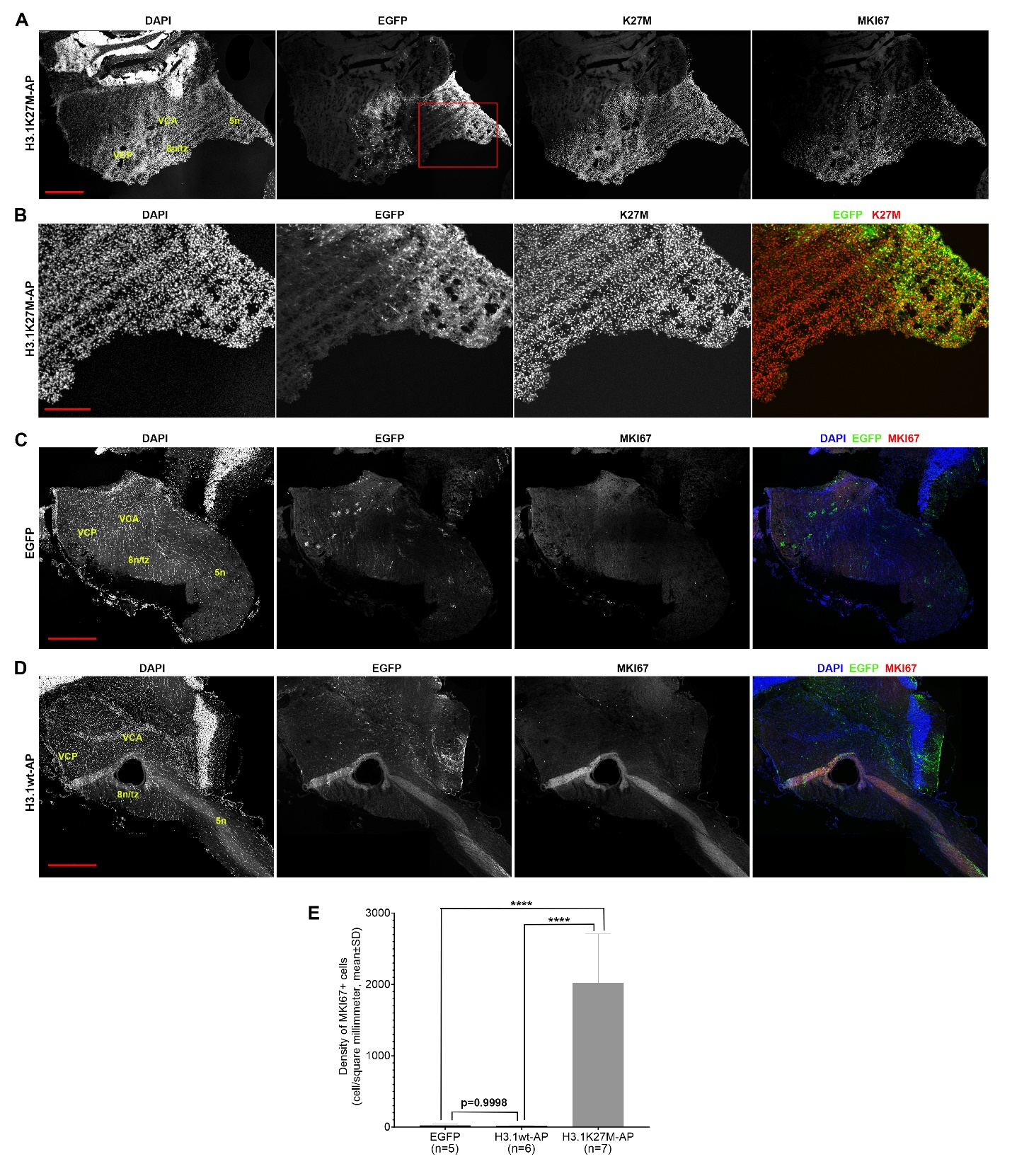

Fig. S4. Co-expression of wild-type H3.1 with activated ACVR1 and PIK3CA does not induce glioma-like growth in the mouse brainstem. (A) Not all transfected cells can be identified by EGFP in mice born from electroporated embryos. Composite confocal images of a P43 mouse in the H3.1K27M-AP group showing that many cells in this sample expressed the H3.1-K27M protein but not EGFP. The boxed area in (A) is shown in (B) at a higher magnification. Note that in (B) most tumor cells on the left side of the panel expressed H3.1-K27M but not EGFP. (C and D) Few proliferating cells are present in the brainstem of mice expressing EGFP alone or EGFP together with wild-type H3.1, ACVR1-G356D, and PIK3CA-E545K (i.e. H3.1wt-AP). Composite images of a P135 mouse of the EGFP group (C) and a P74 mouse of the H3.1wt-AP group (D) are shown. (E) Quantification of the number of proliferating cells in transfected areas of the brainstem. Both EGFP^+^ cells and EGFP^-^ cells in transfected areas were counted. ****p<0.0001, one-way ANOVA. The n values represent the number of embryos used for quantification in each group. 5n: trigeminal nerve; 8n: the 8th cranial nerve; tz: trapezoid body; VCA: ventral cochlear nucleus, anterior; VCP: ventral cochlear nucleus, posterior. Scale bars: 500μm in (A), (C), and (D), 200μm in (B).

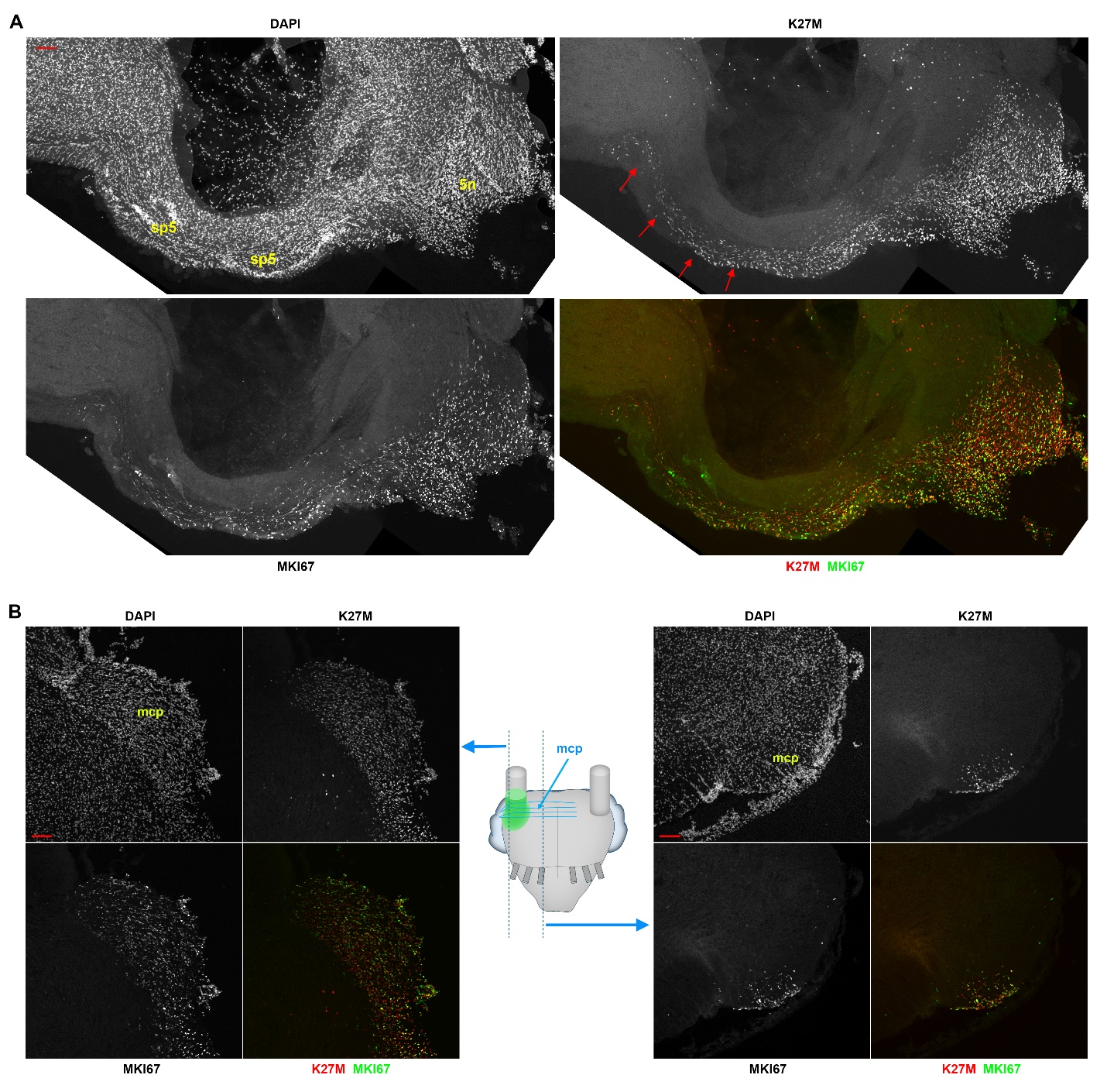

Fig. S5. The presence of proliferating cells expressing H3.1K27M-AP on axon tracts away from but directly linked to TREZ. (A) Composite confocal images of the brainstem of a P22 mouse expressing H3.1K27M-AP. Arrows indicate the part of the spinal trigeminal tract (sp5) relatively far away from the entry point of the trigeminal nerve (5n). (B) Confocal images of the middle cerebellar peduncle (mcp) area in the brainstem of a P29 mouse expressing H3.1K27M-AP. Left panels show mcp at lateral pons where it intersects with the trigeminal nerve, right panels show mcp at more medial side of the pons. Images are representative of eight mice expressing H3.1K27M-AP for both (A) and (B). Scale bars: 100μm.

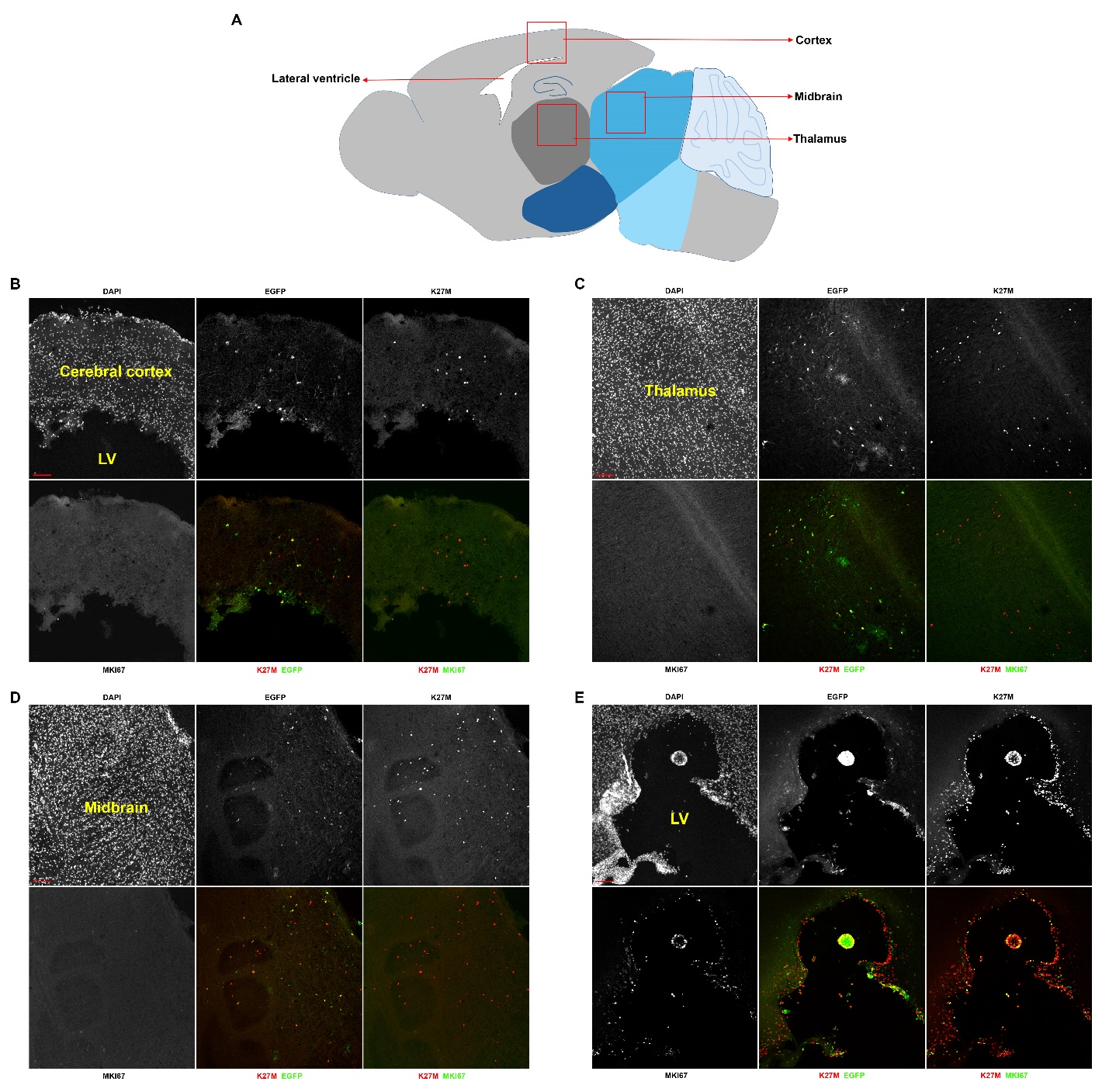

Fig. S6. Anatomical specificity of H3.1K27M-AP induced glioma cell proliferation. (A) Schematic representation of the typical transfected regions in mice that had been subjected to forebrain/midbrain in utero electroporation at E11.5. (B-E) H3.1K27M-AP does not induce glioma-like growth in the parenchyma of the cerebral cortex (B), in the thalamus (C), or in the midbrain (D), but induces periventricular/ventricular gliomagenesis in a subset of mice (E). Images are representative of 13 mice (P21-P135) for the cerebral cortex group, six mice (P21-P47) for the thalamus group, five mice (P25-P69) for the midbrain group, and two mice (P73 and P130; out of the 13 forebrain transfected mice) for the periventricular/ventricular gliomagenesis group. LV: lateral ventricle. Scale bars: 100μm.

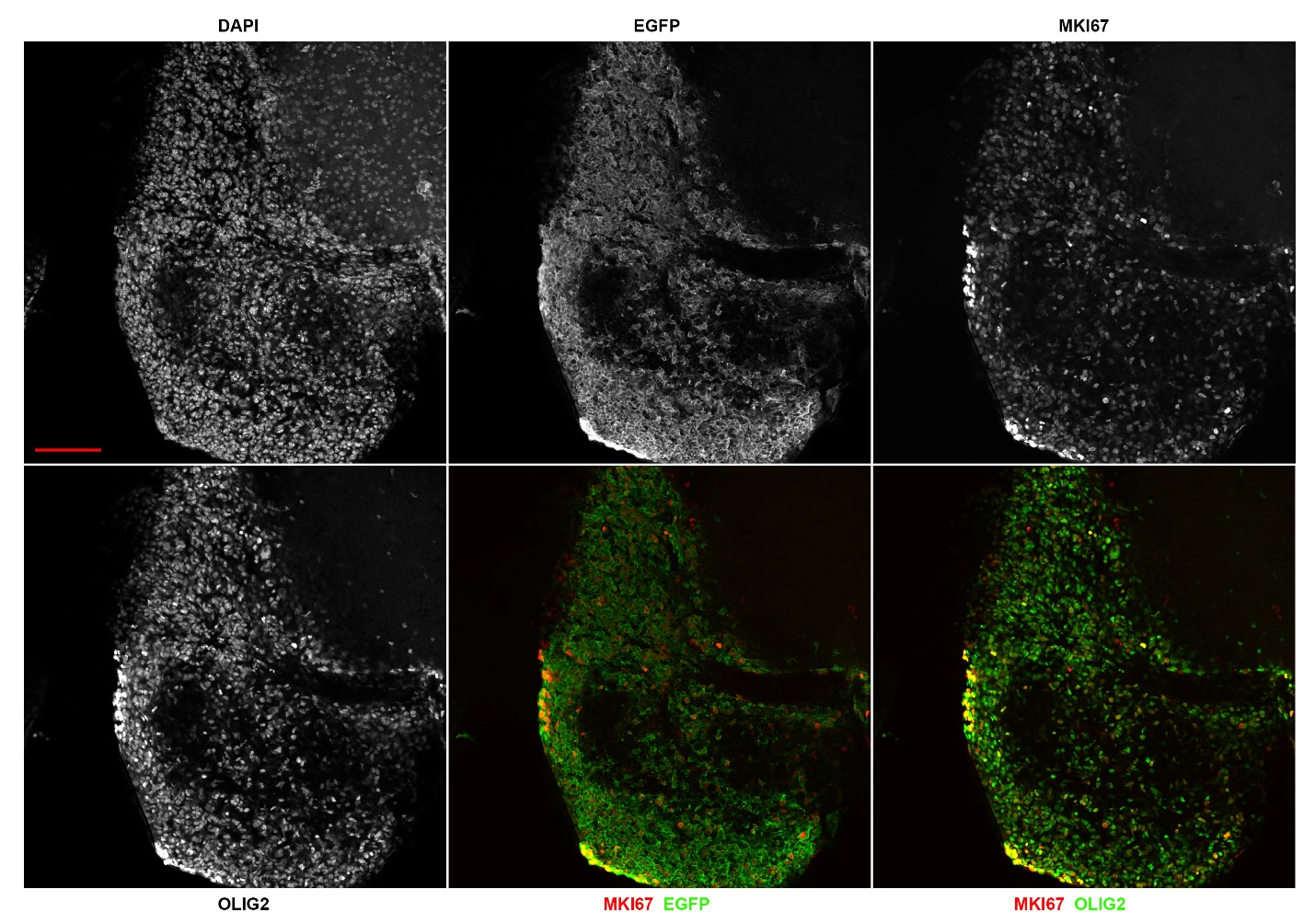

Fig. S7. Expression of OLIG2 in periventricular/ventricular gliomas induced by H3.1K27M-AP. Confocal images of tumor cells at the periventricular/ventricular area from a P73 mouse are shown. Nearly all proliferating cells in the tumor tissue express OLIG2. Images are representative of two mice in which forebrain expression of H3.1K27M-AP induced periventricular/ventricular tumor growth. Scale bar: 100μm.

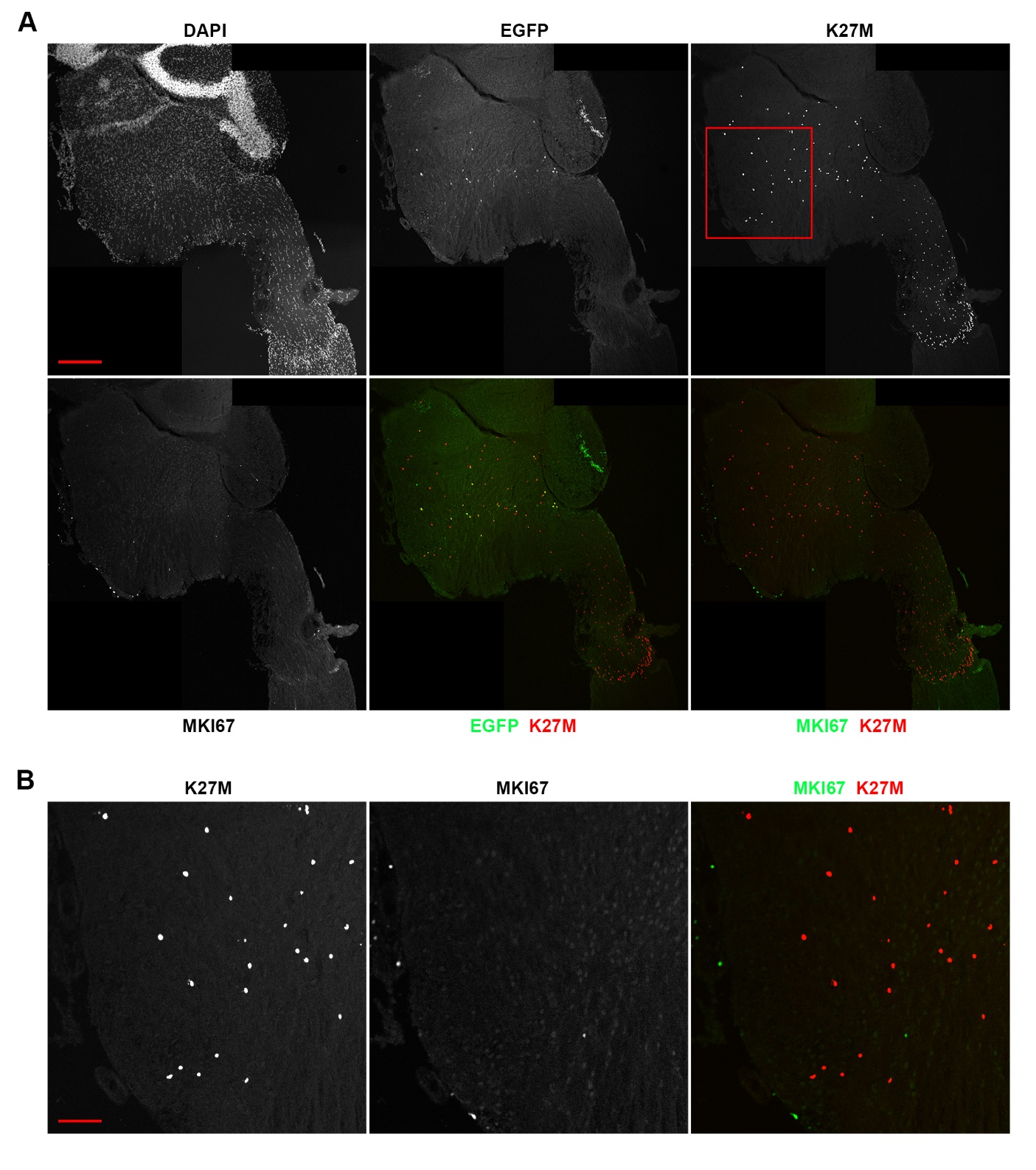

**Fig. S8. Moderate expression of *H3.3K27M* together with its common co-occurring DIPG mutations does not induce cell proliferation in the mouse brainstem.** (**A**) Composite confocal images of the brainstem of a P52 mouse expressing H3.3-K27M, TRP53-R172H, PDGFRA, and CCND2 (H3.3K27M-TPC). EGFP was detected in few cells within the brainstem, and K27M immunoreactivity revealed more but still moderate number of cells within the brainstem. Boxed area in (A) is shown in (**B**) at higher magnification. Note that most of the scattered proliferating cells in the brainstem are not transfected cells as they do not express K27M. Images are representative of seven mice at the age of P45-P183 transfected with H3.3K27M-TPC at moderate efficiencies. Scale bars: 300μm in (A) and 100μm in (B).

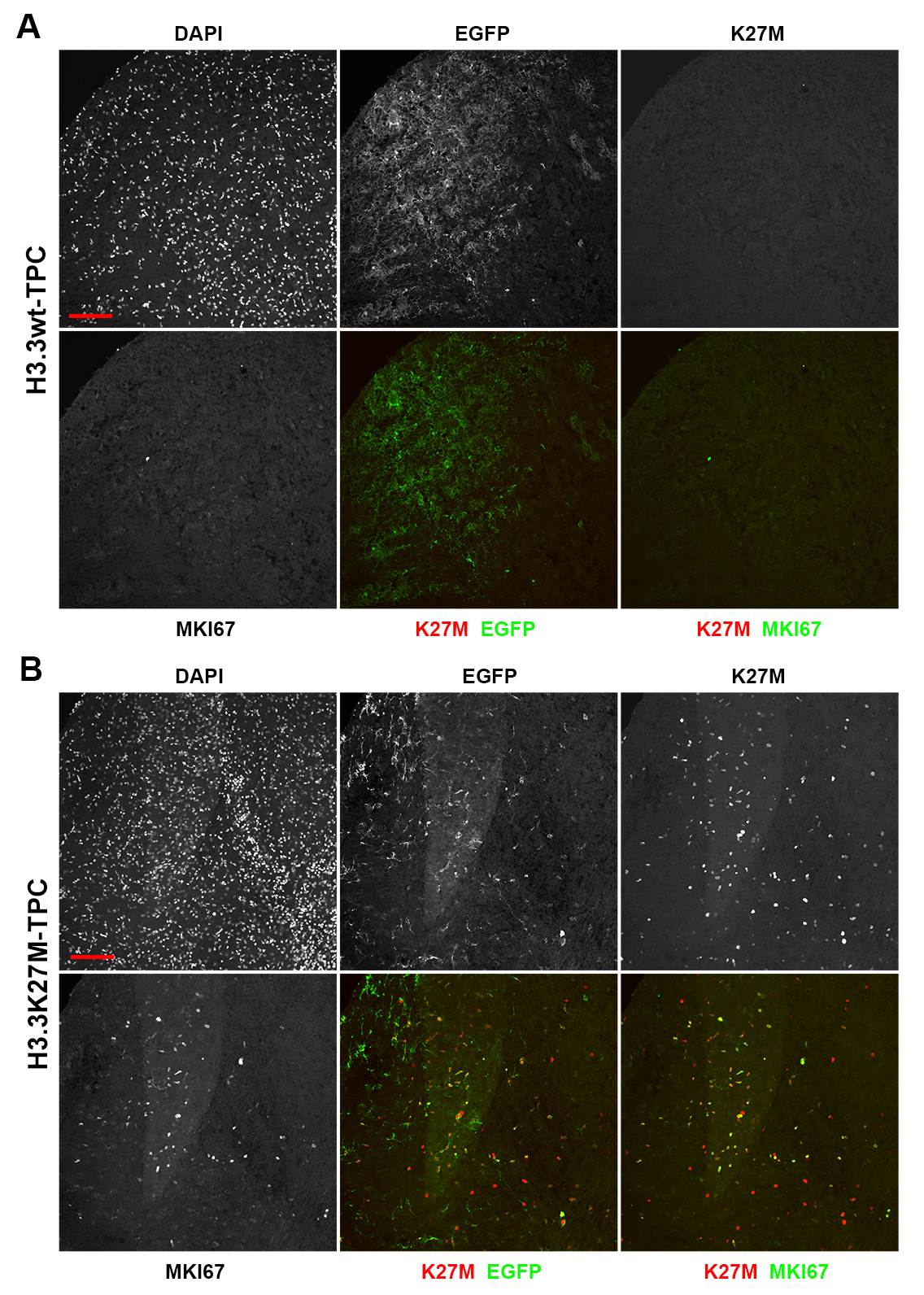

**Fig. S9. High expression of *H3.3K27M* together with its common co-occurring DIPG mutations leads to tumor-like growth in the brainstem.** Transposon-based plasmids for expressing H3.3-K27M, TRP53-R172H, PDGFRA, and CCND2 (H3.3K27M-TPC) were introduced into the mouse brainstem via in utero electroporation at E11.5. In control experiments (the H3.3wt-TPC group), the plasmid for expressing H3.3-K27M was replaced with a plasmid for expressing wild-type H3.3. Confocal images shown are the cochlear nucleus areas from a P98 mouse of the control group and a P125 mouse of the H3.3K27M-TPC group. No K27M immunoreactivity and few proliferating cells were detected in the control section, whereas extensive proliferating K27M^+^ cells were detected in the H3.3K27M-TPC section. Images are representative of seven control mice with high transfection efficiencies at the age of P81-P165 and 10 mice of the H3.3K27M-TPC group with high transfection efficiencies at the age of P47-P163. Scale bars: 100μm.

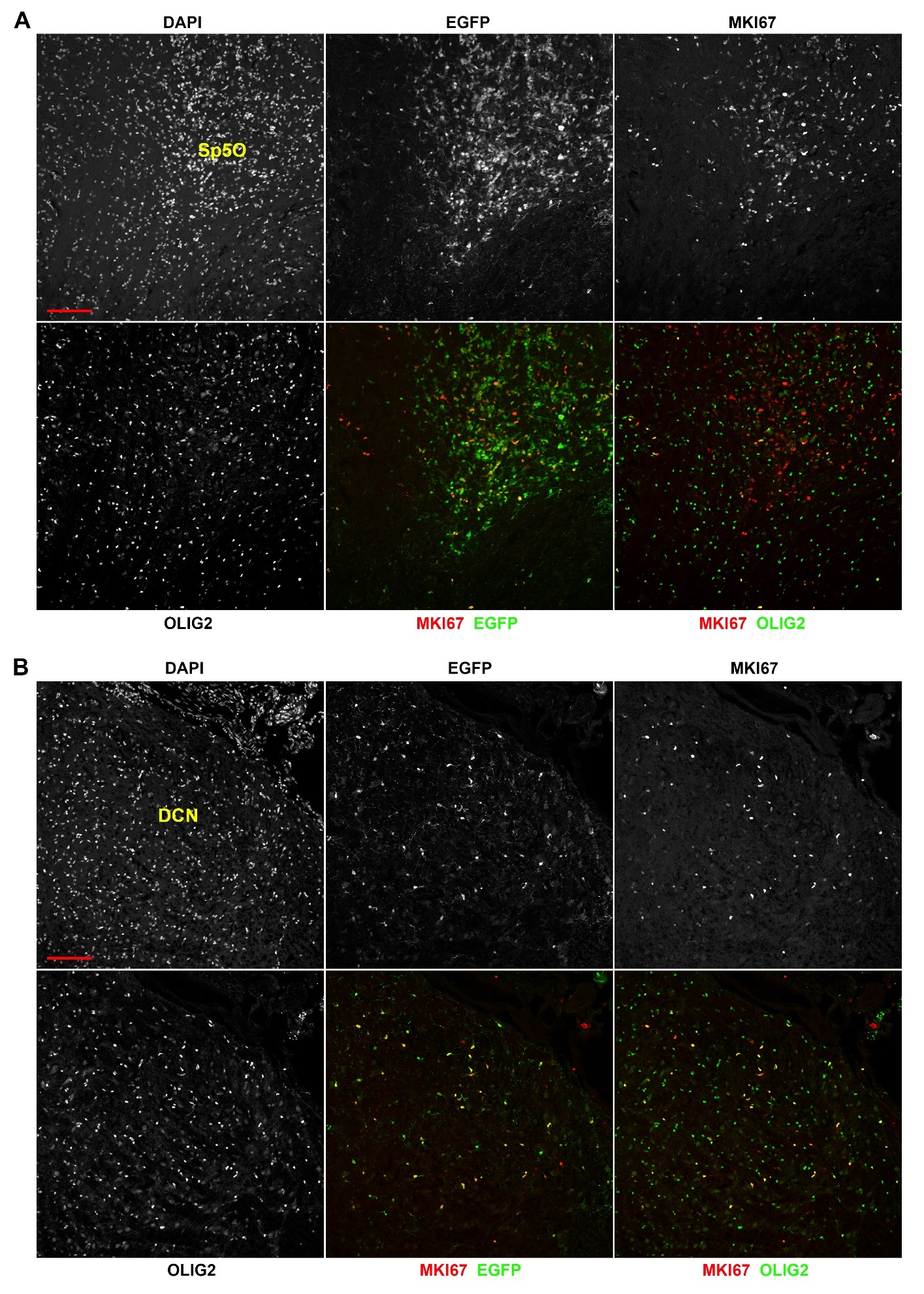

Fig. S10. Tumor-like growth induced by co-expression of *H3.3K27M* and its common co-occurring DIPG mutations include both OLIG2^+^ and OLIG2^-^ cell populations. Confocal images of proliferating cells in the brainstem of a P163 mouse expressing H3.3K27M-TPC. (A) An example of a cluster of proliferating cells that includes both OLIG2^+^ and OLIG2^-^ cells. (B) An example of a cluster of proliferating cells that are mostly OLIG2^+^. DCN: dorsal cochlear nucleus; Sp5O: spinal trigeminal nucleus, oral part. Scale bars: 100μm.

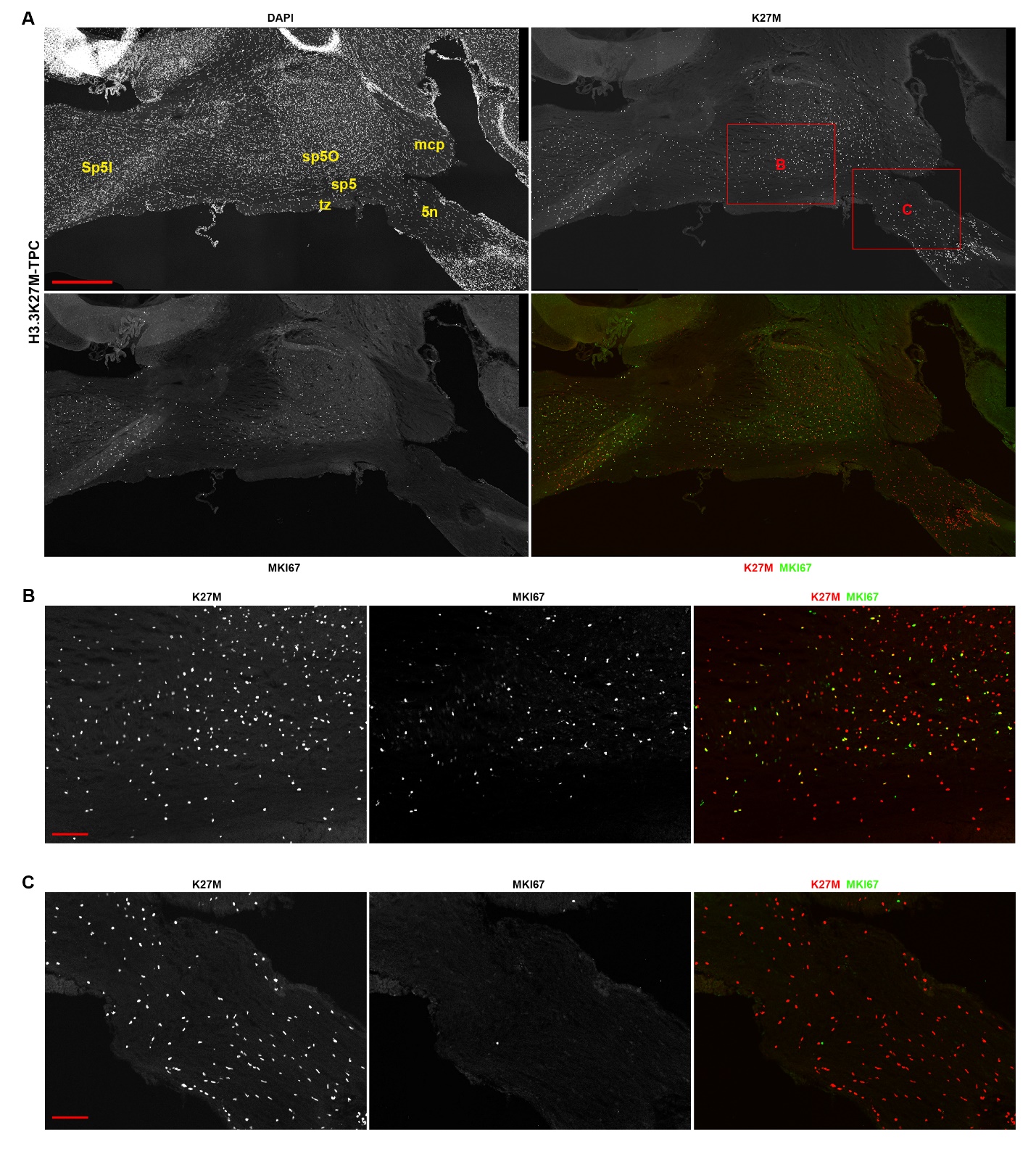

Fig. S11. Glioma-like growth induced by *H3.3K27M* and its common co-occurring DIPG mutations occurs outside of TREZ. (A) Composite confocal images of the brainstem of a P163 mouse expressing H3.3K27M-TPC. Proliferating cells are distributed extensively in both the pons and the medulla. Boxed areas in (A) are shown in (B) (area outside of TREZ) and (C) (area at TREZ) at higher magnifications to illustrate that extensive proliferation of K27M^+^ cells occurs outside of TREZ but not inside TREZ. Images are representative of 10 mice (P47-P163) transfected with H3.3K27M-TPC at high efficiencies. 5n: trigeminal nerve; mcp: middle cerebellar peduncle; sp5: spinal trigeminal tract; Sp5O: spinal trigeminal nucleus, oral part; Sp5I: spinal trigeminal nucleus, interpolar; tz: trapezoid body. Scale bars: 500μm in (A), 100μm in (B) and (C).

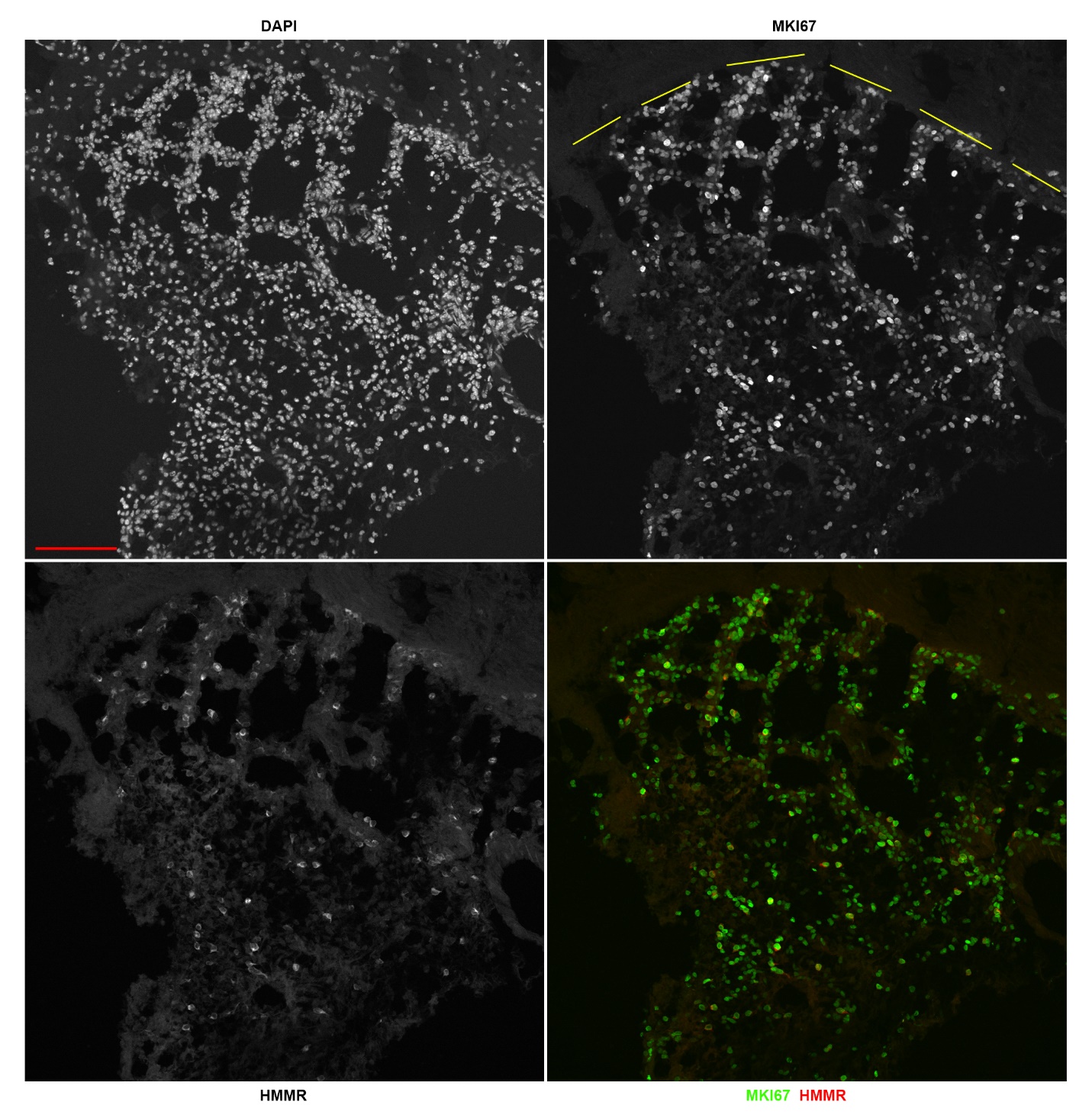

Fig. S12. HMMR expression is increased in gliomas induced by H3.1K27M-AP. Confocal images of the tumor tissue expressing H3.1K27M-AP from a P53 mouse showing expression of HMMR. Dashed lines approximately outline the edge of the tumor tissue. Levels of HMMR in tumor cells are higher compared to nearby non-tumor cells. Images are representative of nine mice in which H3.1K27M-AP-induced large tumor bulk has been formed. Scale bars: 100μm.

Table S1. Mice used in the in utero electroporation analysis

| Group | Sex | Age | Location of IUE | Hydrocephalus (H) or small body size (S) | Head tilt (HT) | Glioma/glioma-like growth (G) |
| --- | --- | --- | --- | --- | --- | --- |
| EGFP | M | P30 | brainstem |  |  |  |
| EGFP | F | P34 | brainstem | H |  |  |
| EGFP | M | P64 | brainstem |  |  |  |
| EGFP | M | P91 | brainstem |  |  |  |
| EGFP | F | P135 | brainstem |  |  |  |
| EGFP | M | P135 | brainstem |  |  |  |
| H3.1wt-AP | F | P60 | brainstem |  |  |  |
| H3.1wt-AP | F | P74 | brainstem |  |  |  |
| H3.1wt-AP | M | P74 | brainstem | H |  |  |
| H3.1wt-AP | M | P75 | brainstem |  |  |  |
| H3.1wt-AP | F | P90 | brainstem |  |  |  |
| H3.1wt-AP | M | P90 | brainstem | H |  |  |
| H3.1wt-AP | M | P159 | brainstem |  |  |  |
| H3.1wt-AP | M | P159 | brainstem |  |  |  |
| H3.1wt-AP | F | P159 | brainstem |  |  |  |
| H3.1wt-AP | F | P159 | brainstem |  |  |  |
| H3.1K27M-AP | M | P40 | brainstem | H |  | G |
| H3.1K27M-AP | F | P42 | brainstem |  | HT | G |
| H3.1K27M-AP | F | P43 | brainstem |  | HT | G |
| H3.1K27M-AP | F | P49 | brainstem |  | HT | G |
| H3.1K27M-AP | F | P52 | brainstem |  | HT | G |
| H3.1K27M-AP | F | P53 | brainstem |  | HT | G |
| H3.1K27M-AP | F | P63 | brainstem |  | HT | G |
| H3.1K27M-AP | M | P64 | brainstem |  | HT | G |
| H3.1K27M-AP | M | P90 | brainstem |  |  | G |
| H3.1K27M-AP | M | P20 | brainstem | H |  | G* |
| H3.1K27M-AP | F | P21 | brainstem | H |  | G* |
| H3.1K27M-AP | F | P21 | brainstem | H |  | G* |
| H3.1K27M-AP | M | P21 | brainstem |  |  | G* |
| H3.1K27M-AP | M | P22 | brainstem | H |  | G* |
| H3.1K27M-AP | F | P29 | brainstem | H |  | G* |
| H3.1K27M-AP | M | P32 | brainstem | H |  | G* |
| H3.1K27M-AP | M | P41 | brainstem | H |  | G* |
| H3.1K27M-AP | M | P56 | brainstem | H |  | G* |
| H3.1K27M-AP | F | P63 | brainstem |  |  | G* |
| H3.1K27M-AP | F | P91 | brainstem |  |  | G* |
| H3.1K27M-AP, HMMR | F | P28 | brainstem | H |  | G |
| H3.1K27M-AP, HMMR | F | P60 | brainstem |  |  | G |
| H3.1K27M-AP, HMMR | M | P61 | brainstem | H | HT | G |
| H3.1K27M-AP, DN-HMMR | F | P21 | brainstem | S |  | G** |
| H3.1K27M-AP, DN-HMMR | M | P22 | brainstem | S |  | G** |
| H3.1K27M-AP, DN-HMMR | M | P22 | brainstem | S |  | G** |
| H3.1K27M-AP, DN-HMMR | M | P53 | brainstem | H | HT | G** |
| H3.1K27M-AP, DN-HMMR | M | P55 | brainstem |  | HT | G** |
| H3.1K27M-AP, DN-HMMR | F | P60 | brainstem | H |  | G** |
| H3.1K27M-AP, DN-HMMR | M | P60 | brainstem |  |  | G** |
| H3.1wt-AP | F | P19 | FB/MB | H |  |  |
| H3.1wt-AP | F | P21 | FB/MB | H |  |  |
| H3.1wt-AP | F | P22 | FB/MB | H |  |  |
| H3.1wt-AP | M | P27 | FB/MB | H |  |  |
| H3.1wt-AP | F | P28 | FB/MB | H |  |  |
| H3.1wt-AP | F | P60 | FB/MB |  |  |  |
| H3.1wt-AP | M | P60 | FB/MB |  |  |  |
| H3.1wt-AP | F | P91 | FB/MB |  |  |  |
| H3.1wt-AP | F | P91 | FB/MB |  |  |  |
| H3.1K27M-AP | M | P21 | FB/MB | H |  |  |
| H3.1K27M-AP | M | P21 | FB/MB |  |  |  |
| H3.1K27M-AP | M | P23 | FB/MB | H |  |  |
| H3.1K27M-AP | F | P25 | FB/MB |  |  |  |
| H3.1K27M-AP | M | P25 | FB/MB |  |  |  |
| H3.1K27M-AP | M | P25 | FB/MB |  |  |  |
| H3.1K27M-AP | M | P25 | FB/MB |  |  |  |
| H3.1K27M-AP | M | P28 | FB/MB | H |  |  |
| H3.1K27M-AP | M | P29 | FB/MB | H |  |  |
| H3.1K27M-AP | M | P31 | FB/MB | H |  |  |
| H3.1K27M-AP | F | P47 | FB/MB | H |  |  |
| H3.1K27M-AP | M | P69 | FB/MB | H |  |  |
| H3.1K27M-AP | F | P73 | FB/MB | H |  | G*** |
| H3.1K27M-AP | M | P130 | FB/MB | H |  | G*** |
| H3.1K27M-AP | M | P135 | FB/MB |  |  |  |
| H3.1K27M-AP | F | P135 | FB/MB |  |  |  |
| H3.3wt-TPC | M | P81 | brainstem | H |  |  |
| H3.3wt-TPC | M | P98 | brainstem | H |  |  |
| H3.3wt-TPC | M | P98 | brainstem |  |  |  |
| H3.3wt-TPC | F | P98 | brainstem |  |  |  |
| H3.3wt-TPC | F | P98 | brainstem |  |  |  |
| H3.3wt-TPC | M | P165 | brainstem |  |  |  |
| H3.3wt-TPC | M | P165 | brainstem |  |  |  |
| H3.3K27M-TPC | M | P45 | brainstem | H |  |  |
| H3.3K27M-TPC | F | P47 | brainstem | H |  | G* |
| H3.3K27M-TPC | F | P48 | brainstem | H |  | G* |
| H3.3K27M-TPC | F | P48 | brainstem |  |  |  |
| H3.3K27M-TPC | F | P49 | brainstem |  |  | G* |
| H3.3K27M-TPC | F | P52 | brainstem |  |  |  |
| H3.3K27M-TPC | F | P52 | brainstem |  |  | G* |
| H3.3K27M-TPC | M | P52 | brainstem | H |  | G* |
| H3.3K27M-TPC | M | P54 | brainstem | H |  |  |
| H3.3K27M-TPC | M | P97 | brainstem | H |  | G* |
| H3.3K27M-TPC | M | P125 | brainstem | H |  | G* |
| H3.3K27M-TPC | F | P163 | brainstem |  |  |  |
| H3.3K27M-TPC | F | P163 | brainstem |  |  | G* |
| H3.3K27M-TPC | F | P163 | brainstem |  |  | G* |
| H3.3K27M-TPC | M | P163 | brainstem |  |  |  |
| H3.3K27M-TPC | M | P163 | brainstem |  |  | G* |
| H3.3K27M-TPC | F | P183 | brainstem |  |  |  |

G*: Extensive glioma-like cell proliferation without apparent large tumor bulk.

G**: Proliferating tumor cells were those that appeared to have lost the expression of DN-HMMR.

G***: Ventricular/periventricular gliomagenesis.

FB/MB: Forebrain/midbrain

Movie S1.

Head-tilting phenotype of a P43 female mouse transfected with H3.1K27M-AP.
